# Amyloid Beta Oligomers Accelerate ATP-Dependent Phase Separation of miRNA-Bound Ago2 to RNA Processing Bodies *in vitro*

**DOI:** 10.1101/2024.03.14.584939

**Authors:** Sritama Ray, Sumangal Roychowdhury, Yogaditya Chakraborty, Saikat Banerjee, Krishnananda Chattopadhyay, Kamalika Mukherjee, Suvendra N. Bhattacharyya

## Abstract

Phase separation to insoluble membrane-less organelles is a major way of activity regulation of specific proteins in eukaryotic cells. miRNA-repressed mRNAs and Ago proteins are known to be localized to RNA-processing bodies, the subcellular structures which are formed due to assembly of several RNA binding and regulatory proteins in eukaryotic cells. Ago2 is the most important miRNA binding protein that by forming complex with miRNA binds to mRNAs having cognate miRNA binding sites and represses protein synthesis in mammalian cells. Factors which control compartmentalization of Ago2 and miRNA-repressed mRNAs to RNA processing bodies are largely unknown. We have adopted a detergent permeabilized cell-based assay system to follow the phase separation of exogenously added Ago2 to RNA processing bodies *in vitro*. The Ago2 phase separation process is ATP dependent and is influenced by osmolarity and salt concentration of the reaction buffer. miRNA binding of Ago2 is essential for its targeting to RNA processing bodies and the compartmentalization process gets retarded by miRNA binding “sponge” protein HuR. This assay system found to be useful in identification of amyloid beta oligomers as miRNA-activity modulators which repress miRNA activity by enhancing Ago2-miRNP targeting to RNA processing bodies.

**Graphical Abstract:** 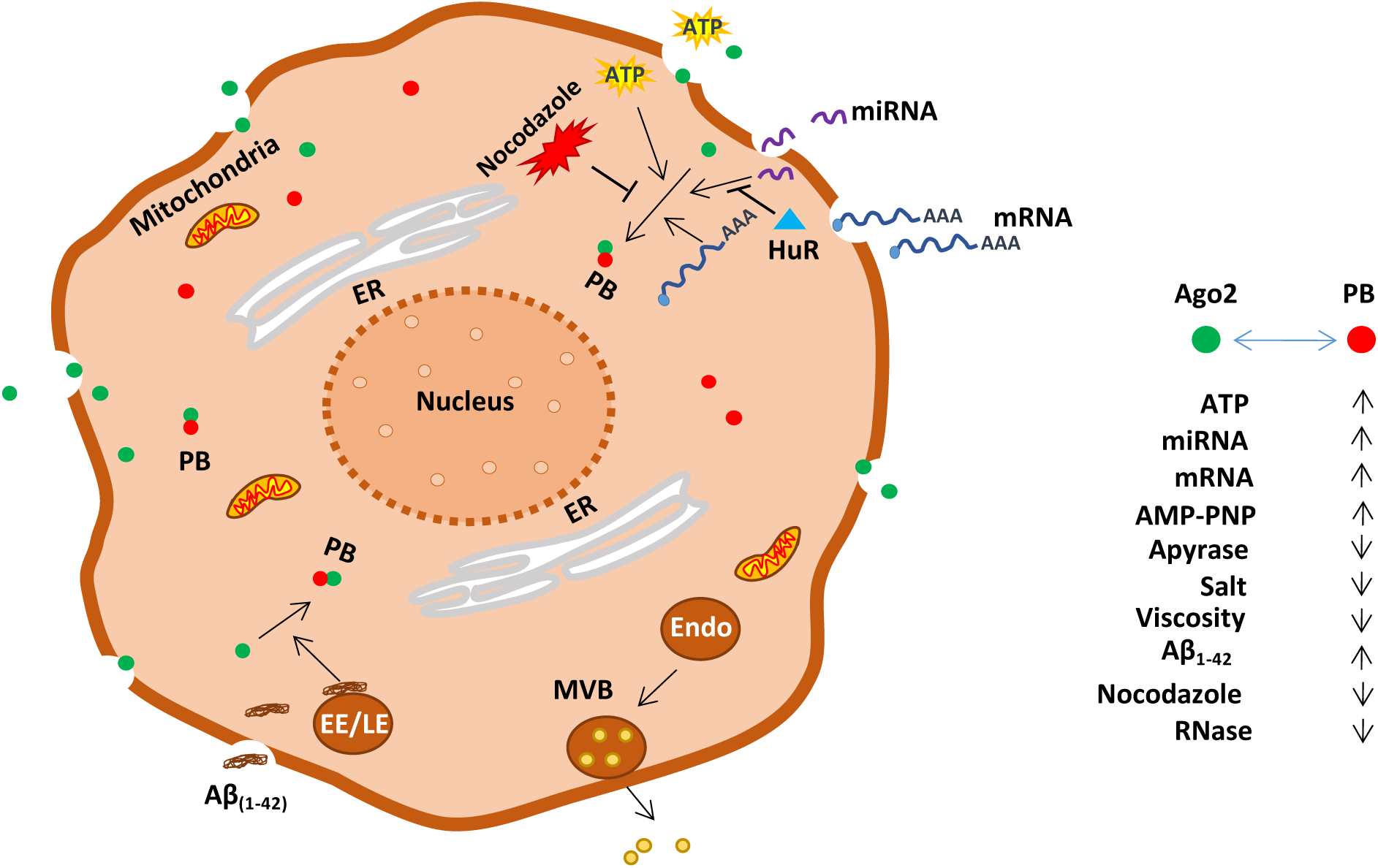

- miRNA bound Ago2 gets phase separated *in vitro* to RNA processing bodies (PBs) in detergent permeabilized mammalian cells.
- Phase separation of Ago2 to PBs is controlled by presence of ATP and RNA.
- Amyloid beta oligomers retard dynamics of Ago2 bodies to inhibit miRNA function and enhance PB targeting of Ago2 miRNPs.
- microRNA binding protein HuR can rescue Ago2 miRNP from PBs and inverse the effect of amyloid beta oligomers.

## Introduction

miRNAs are 22 nucleotides long regulatory RNA expressed by metazoan. The mature miRNA by forming complex with Argonaute (Ago) proteins binds target mRNAs with perfect or imperfect complementarities to repress protein expression(1). miRNAs primarily interact with the target messages via complementary binding sites on the 3’ UTR of mRNAs(2). Interestingly, the fate of miRNP-bound target messages depends on the degree of complementarity in miRNA-mRNA interaction. While a fully complementary binding promotes the endo nucleolytic activity of Ago2 protein leading to the degradation of the target messages(3), the imperfect complementarity allows the target messages to either get degraded or to remain stored in the translationally inactive state in specialized subcellular structures called RNA-processing bodies or PBs(4). The miRNA-Ago2 complex, the miRISC, remains localized to PBs along with repressed mRNAs in diverse types of mammalian cells including mammalian neurons (5,6). Interestingly, several components of PBs are known to have specific roles in RNA degradation. PBs are mainly composed of many proteins involved in translational repression or degradation of mRNAs such as deadenylation complex Lsm 1-7, Ccr4-Not, decapping co-activator and enzyme Dcp1/Dcp2, decapping activators Edc3, DDX6, 5’ to 3’ exoribonuclease Xrn1(7) and mRNA binding proteins like 4E-T, CPEB1 which promote translational repression (8). Ago2 interacting GW182 proteins also accumulate into PBs and are responsible for stability of miRNAs in mammalian cells. They, possibly by facilitating Ago2 miRNPs localization to PBs, prevent export of “used” miRNPs from mammalian cells(9). Interestingly, while a group of miRNA-mRNA complexes get targeted to P-bodies possibly for subsequent degradation, a subset of miRNA-repressed mRNAs escapes the degradation path and remains stored in PBs and are reused for protein translation during altered physiological context like refeeding of amino acid-starved hepatic cells (10). In more recent time, purification and *in situ* analysis of PBs confirmed species-specific compartmentalization of specific mRNAs to PBs(11). Therefore, the mRNA targeting and relocalization from PBs are two important events contributing to cellular physiology and stress management processes(12). Unfortunately, the mechanism of PB-compartmentalization of mRNAs or Ago proteins are not well studied due to lack of *in vitro* assay system to follow the same.

Liquid-liquid phase separation (LLPS) has emerged recently as a vital mechanism explaining formation and function of various membrane less structures present in the cell. Macromolecules undergoing LLPS tend to condense into a dense phase which resembles liquid droplets like structures, co-existing with surrounding dilute phase (13). In recent studies, LLPS has emerged as key mechanism regulating a wide variety of biological activities including adaptive and innate immune signaling, transcription, autophagy and PB assembly (14). Any aberration in LLPS processes is expected to develop dysregulated gene expression resulting in different disease conditions. Protein aggregation is associated with neurodegenerative processes.

Previous report suggests Ago2 phase separation to PBs as a dynamic event that is responsive to neuronal stimulation in nerve cells(15). Therefore, any study that can reveal the parameters of this dynamic process of Ago, miRNA or target mRNA phase separation will not only allow us to understand the biochemical properties of this event but also will help us to develop chemical modifiers/inhibitors to affect Ago2 phase separation that will alter miRNA activity in neuronal and non-neuronal cells to control diseases attributed by LLPS of Ago2 or related proteins.

*In vitro* phase separation study of Ago proteins to PBs has not been achieved before. One major challenge was related to having purified PBs from mammalian cells without altered biochemical compositions. We have developed an assay system using the detergent (Digitonin) permeabilized HeLa cells to follow the trafficking of Ago2 proteins added exogenously to the permeabilized cells. The added Ago2 gets phase separated to form bodies which colocalizes with PBs present in permeabilized HeLa cells. The compartmentalization of Ago2 to PBs is ATP dependent and gets retarded by RNase treatment. Both osmolarity and viscosity of the reaction buffer have effects on Ago2 phase separation to PBs. As reported in the *in vivo* conditions, the miRNA and miRNA-targeted mRNAs were also detected in PBs in the *in vitro reaction*. The miRNA binding to Ago2 is necessary for Ago2 localization to PBs and miRNA binding protein HuR retarded both miRNA and Ago2 localization to PBs. Interestingly, the arbitration of the Ago2 localization can also be achieved by Nocodazole, an agent that affect tubulin cytoskeleton dynamics. Therefore, the assay system described here can be useful to identify agents that, by affecting Ago2/miRNA targeting to RNA processing bodies, can alter gene repression by miRNAs in higher eukaryotic cells. Interestingly, amyloidogenic amyloid beta oligomers could control the PB compartmentalization of Ago2 by promoting the phase separation of Ago2 to reduce the miRNA activity in cells exposed to Aβ_(1-42)_ oligomers *ex vivo*. This is consistent with the *in vivo* data in AD brain (16,17).

## Results

### Phase separated Ago2 bodies are formed *in vitro* by exogenous protein in detergent-permeabilized mammalian cells

We were curious to study the phase separation process of Ago2 protein that is related to miRNA activity loss and large Ago2 body formation in differentiating neuronal cells (15). Inhibition of protein translation process in mammalian cells can affect Ago2 body formation also (18). To mechanistically understand the process of how Ago2 bodies are formed, we found it essential to have an *in vitro* assay system to study the dynamicity and biochemical requirement of Ago2 body formation process. The cellular scaffold structure may also be essential to form Ago2 bodies that are generated from the assembly of soluble Ago2 protein present in the reaction buffer. We have used HeLa cells to get it permeabilized with the mild detergent Digitonin to allow passage of exogenously added Ago2 protein through the porous membrane to reach cytoplasm, interact with cellular structures and form bodies. To distinguish the newly formed bodies from existing Ago2 bodies, we have incubated Digitonin-permeabilized Hela cells with isotonic lysates prepared by sonication of HEK293 cells expressing FH-Ago2. The entry and body-forming ability of the FH-Ago2 was monitored against time at 37°C (Fig. 1A). To standardize the condition we have used increasing concentration of Digitonin to permeabilize the HeLa cells to find 80ng/ml as the optimum concentration that doesn’t affect cell morphology and integrity of existing structures while allowing the internalization of Ago2 and formation of bodies inside the permeabilized recipient HeLa cells (Fig 1B and G). We have incubated HeLa cells, permeabilized with 80 ng/ml digitonin in 1X PBS for 10 minutes, with HEK293 lysate at two different incubation temperatures to observe the effect of temperature on internalization of FH-Ago2 and body formation. The number of Ago2 bodies observed at 37°C was much higher than at 25°C incubation temperature (Figure 1C, D and H). We have tried different time of incubation with FH-Ago2 containing HEK293 lysate at 37°C and observed enhanced accumulation of Ago2 bodies with increasing time points in detergent-permeabilized HeLa cells. However, longer incubation with detergent resulted in deformed morphology of HeLa cells. We found 30 minutes of incubation as the optimum condition where reasonable number of Ago2 bodies were detected without much effect of detergent observed on cell morphology and subcellular structures (Fig. 1E, F and I). We did not observe any detrimental effect of 30 minutes of lysate incubation at 37°C with HeLa cells permeabilized with 80ng/ml Digitonin on the subcellular structures like endoplasmic reticulum, endosomes, or mitochondria in HeLa cells after treatment (Supplementary Figure S1A, B).

**Figure 1.**
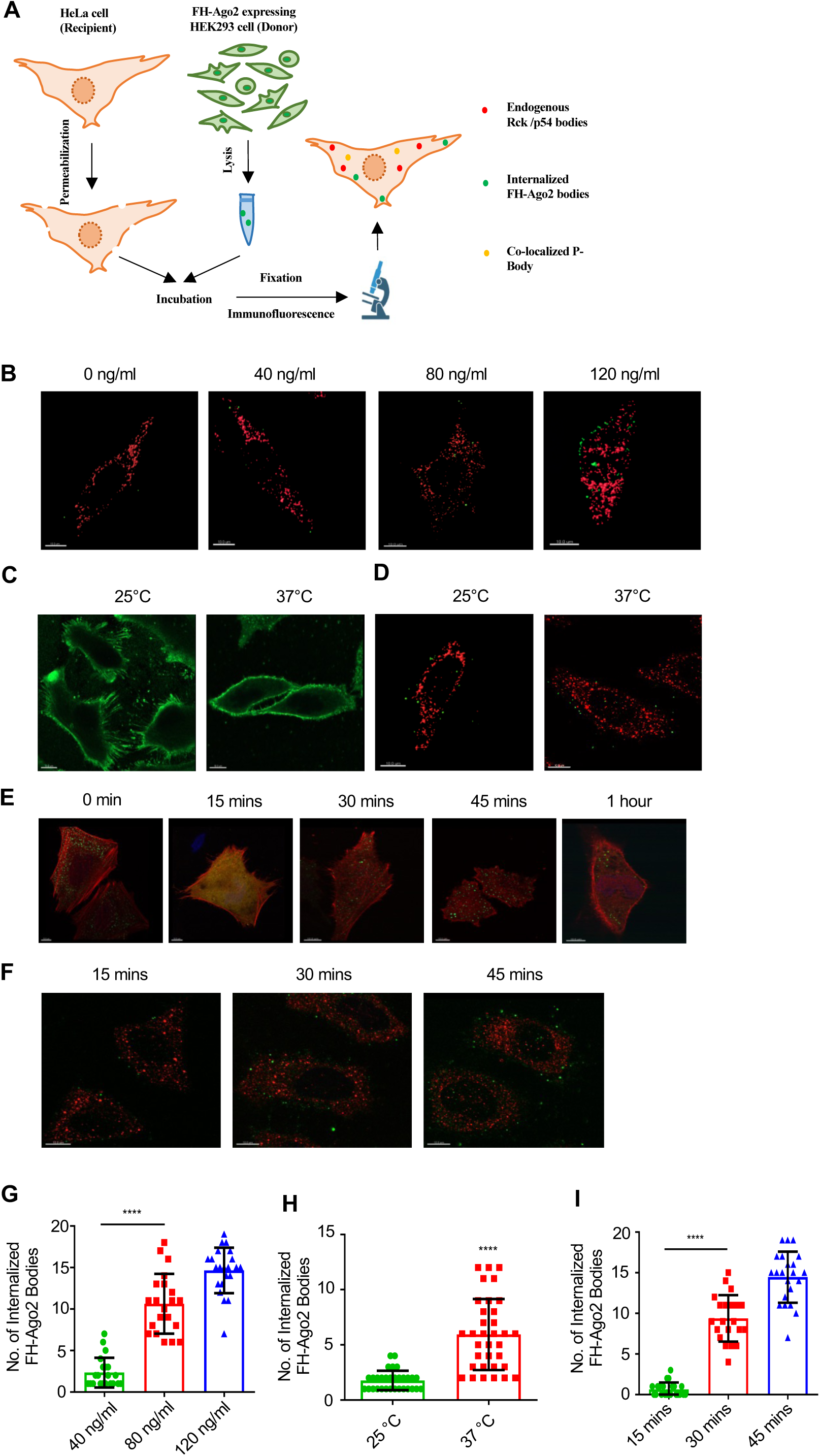
Internalized Ago2 form phase separated bodies in detergent permeabilized HeLa cells. **(A)** A schematic representation of the permeabilization assay using HeLa cells as recipient and HEK293 cells as donor. **(B)**Confocal Images showing internalization of FH-Ago2 proteins (Green) in HeLa cell (recipient) permeabilized with different concentrartion of Digitonin (40 ng, 80 ng, 120 ng), incubated with FH-Ago2 transfected HEK293 cell (donor) lysate and immunostained for endogenous Rck/p54 proteins (Red). **(C)**Confocal Images showing morphology of HeLa cells (recipient) incubated with untransfected HEK293 cell (donor) lysate at different incubation temperature (25 ⁰C and 37 ⁰C) and immunestained for β-Actin (Green). **(D)** Confocal Images showing internalization of FH-Ago2 proteins (Green) in HeLa cell (recipient) incubated with FH-Ago2 transfected HEK293 cell (donor) lysate at different incubation temperature (25 ⁰C and 37 ⁰C) and immunostained for endogenous Rck/p54 proteins (Red).**(E)** Confocal Images showing morphology of HeLa cells (recipient) transfected with Ds Red Actin (Red) and GFP-SMN (green) and incubated with non-transfected HEK293 cell (donor) lysate for different incubation time points (0 minutes, 15 minutes, 30 minutes, 45 minutes, 1 hour). **(F)** Confocal Images showing internalization of FH-Ago2 proteins (Green) in HeLa cell (recipient) incubated with FH-Ago2 transfected HEK293 cell (donor) lysate for different incubation time points (15 minutes, 30 minutes, 45 minutes) and immunostained for endogenous Rck/p54 proteins (Red). **(G)** Quantification of internalized FH-Ago2 proteins (Green) in HeLa cell (recipient) incubated with FH-Ago2 transfected HEK293 cell (donor) lysate when recipient cells were permeabilized with different concentration of Digitonin (40 ng, 80 ng, 120 ng) (p < 0.0001 (between 40 ng and 80 ng groups), n=3, >20 cells used for quantification), (**H**) at different incubation temperature (25 ⁰C and 37 ⁰C) (p < 0.0001, n=3, >30 cells used for quantification), **(I)** incubated with FH-Ago2 transfected HEK293 cell lysate for different incubation time points (15 minutes, 30 minutes, 45 minutes) (p < 0.0001 (between 15 minutes and 30 minutes groups), n=3, >20 cells used for quantification). Merged images are given. Fields have been detected in 63X magnification. Scale bar represents 10 µm. Data represents means ± SDs; ns, non-significant, *p < 0.05, **p < 0.01, ***p < 0.001, ****p < 0.0001. p values were obtained by using two-tailed unpaired Student’s t test.

### Internalized Ago2 gets phase separated and colocalizes with RNA processing bodies *in vitro*

We wanted to see if the Ago2 bodies formed with FH-Ago2 in Digitonin permeabilized HeLa cells do get localized with any specific cellular structures. Interestingly, we saw colocalization of these bodies with existing phase separated structures like RNA-processing bodies. We found complete colocalization of the newly formed Ago2 bodies with pre-existing Ago2 bodies in recipient HeLa cells. However, internalized Ago2 also showed a substantial colocalization with GW182 and RCK/p54bodies (Figure 2A). Its colocalization with Dcp1a bodies and Ago2 bodies found to be more proximal with respect to RCK/p54 bodies while the overlapping intensity of Ago2 and Dcp1a is more substantial in bodies positive for both Dcp1a and exogenous FH-Ago2 (Figure 2A and C and supplementary Figure S1D). Interestingly, when NHA-LacZ was expressed in HEK293 cells and the lysate was used for treatment of permeabilized HeLa cells, we did not see uniform structures like the bodies we observed with FH-Ago2, and we did not find any colocalization of internalized NHA-LacZ with RCK/p54 in HeLa cells (Supplementary Figure S1C). Therefore, there have been PBs positive for internalized FH-Ago2 and these may have formed due to possible phase separation of soluble internalized FH-Ago2 protein. No colocalization of newly formed Ago2 was observed with ribosomal component S3 (Supplementary Figure S1C). While measuring the Pearson’s Coefficient of colocalization, we noted highest colocalization of FH-Ago2 with GFP-Ago2 and least with GW182 in recipient HeLa cells (Figure 2D). Interestingly, when tried for other cell types, we also observed internalization and PB targeting of FH-Ago2 in Digitonin permeabilized human hepatoma cell Huh7 and NGF-differentiated rat pheochromocytoma cell PC12 incubated with HEK293 lysate expressing FH-Ago2 (Supplementary Figure S2A).

**Figure 2.**
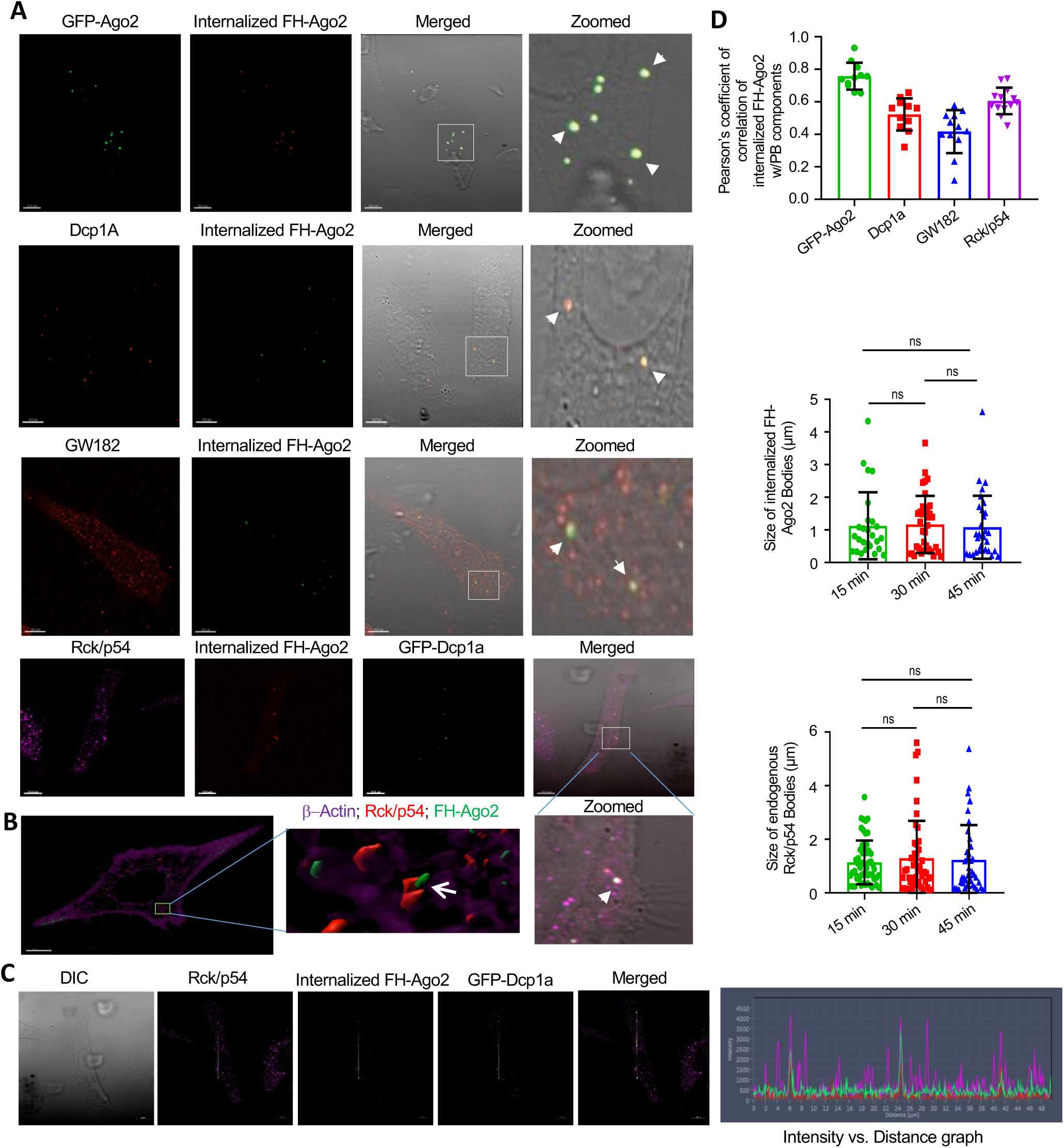
Phase separated Ago2 get localized with RNA processing bodies in detergent permeabilized HeLa cells. **(A)** Confocal Images showing co-localization between internalized FH-Ago2 protein (Red) with exogenously expressed GFP-Ago2 protein (green) in HeLa cells (recipient) incubated with FH-Ago2 transfected HEK293 cell (donor) lysate. Confocal Images showing co-localization between internalized FH-Ago2 protein (Green) and endogenous Dcp1a protein (Red), or endogenous GW182 protein (Red), or exogenously expressed GFP-Dcp1a protein (Green) and endogenous Rck/p54 protein (Violet) in HeLa cells (recipient) incubated with FH-Ago2 transfected HEK293 cell (donor) lysate. **(B)** Z-stack 3D image showing co-localization in 3D between internalized FH-Ago2 protein (Green) and endogenous Rck/p54 protein (Red) in HeLa cells (recipient) incubated with FH-Ago2 transfected HEK293 cell (donor) lysate, β-Actin also got detected by indirect immunofluorescence (Violet). **(C)** Localization and proximity of internalized FH-Ago2 protein (Red), exogenously transfected GFP-Dcp1a protein (Green) and endogenously immunostainedRck/p54 protein (Violet) in HeLa cells (recipient), graph represents intensity profile of all the pixels along the defined section of the cell. Merged images and images obtained by differential interference contrast (DIC) are given. Zoomed images are magnified representation of the inset box in merged images. Fields have been detected in 63X magnification. Scale bar represents 10 µm.**(D)**Pearson’s coefficient of correlation between internalized FH-Ago2 protein (Red for colocalization with GFP-Ago2 and Green for colocalization with Dcp1a/ GW182/ Rck/p54) and different PB components like exogenously expressed GFP-Ago2 (Green), endogenous Dcp1a (Red), GW182 (Red) and Rck/p54 (Violet) in HeLa cell (recipient) incubated with FH-Ago2 transfected HEK293 cell (donor) lysate (upper panel). Size of internalized FH-Ago2 protein (Green) in µm in HeLa cell (recipient) along 15 minutes/ 30 minutes/ 45 minutes time points of HEK293 (donor) cell lysate incubation. (p=0.8682 (between 15 minutes and 30 minutes groups), n=3, >25 cells used for quantification, p=0.7064 (between 30 minutes and 45 minutes groups), n=3, >25 cells used for quantification, p=0.8674 (between 15 minutes and 45minutes groups), n=3, >25 cells used for quantification) (middle panel). Size of endogenous Rck/p54 protein in µm in HeLa cell (recipient) along 15 minutes/ 30 minutes/ 45 minutes time points of HEK293 cell (donor) lysate incubation.(p=0.4782 (between 15 minutes and 30 minutes groups), n=3, >35 cells used for quantification, p=0.8278 (between 30 minutes and 45 minutes groups), n=3, >35 cells used for quantification, p=0.6646 (between 15 minutes and 45minutes groups), n=3, >35 cells used for quantification) (lower panel).Data represents means ± SDs; ns, non-significant, *p < 0.05, **p < 0.01, ***p < 0.001, ****p < 0.0001. p values were obtained by using two-tailed unpaired Student’s t test.

### Effect of altered incubation conditions on PB targeting of Ago2 *in vitro*

To study the effect of changes in different physiochemical properties of the cytoplasm on localization of Ago2 protein to PBs positive for the marker protein Rck/p54, we have altered the cell permeabilization-resealing assay conditions. We have changed the temperature of incubation of recipient HeLa cells with FH-Ago2 expressing HEK293 (donor) cell lysate, to see the effect of temperature on this phase separation process. We have tried 4, 25 and 37°C for incubation and observed that both the internalization (as previously seen during standardization of the assay; Fig 1C, 1D and 1H) and the PB targeting of internalized FH-Ago2 proteins were significantly increased at the physiological temperature 37 °C (Figure 3A and 3B). We were also eager to study the effect of pH of the reaction buffer on PB-targeting of Ago2 and we have changed the pH of the incubation buffer (acidic pH 4.5, physiological pH 7.5 and basic pH 9.0) to see the effect on internalization and PB-targeting of FH-Ago2 in recipient HeLa cell. Association of internalized Ago2 with endogenous PBs has been observed to be the highest at pH 7.5 (Figure 3C and 3D). Recent studies suggest LLPS process also gets affected by the physiochemical property of the solute like osmolarity and viscosity have an prominant effect on LLPS (13). We had increased the salt concentration (200 and 500mM of KCl) of the lysis buffer used for the donor HEK293 cell against the control (78 mM of KCl) condition used in all other assays. We observed that the compartmentalization of internalized FH-Ago2 proteins to endogenous PBs was highly affected by increase in the salt concentration (Figure 3E and 3F). The salt concentration doesn’t have any effect on the stability of FH-Ago2 protein in solution and no change in FH-Ago2 level in lysate was observed when incubated with increasing salt concentration (Figure 3E). We had similar observation when we increased the viscosity of the lysis buffer of donor HEK293 cell by adding 10 and 25 % of Glycerol. There also, internalization as well as targeting of FH-Ago2 protein to the endogenous PBs were significantly decreased with increasing viscosity of the reaction buffer. Interestingly, the donor cell derived FH-Ago2 protein was observed to remain clustered near the cell membrane that failed to get targeted to perinuclear PBs in presence of high glycerol (Figure 3G and 3H).

**Figure 3.**
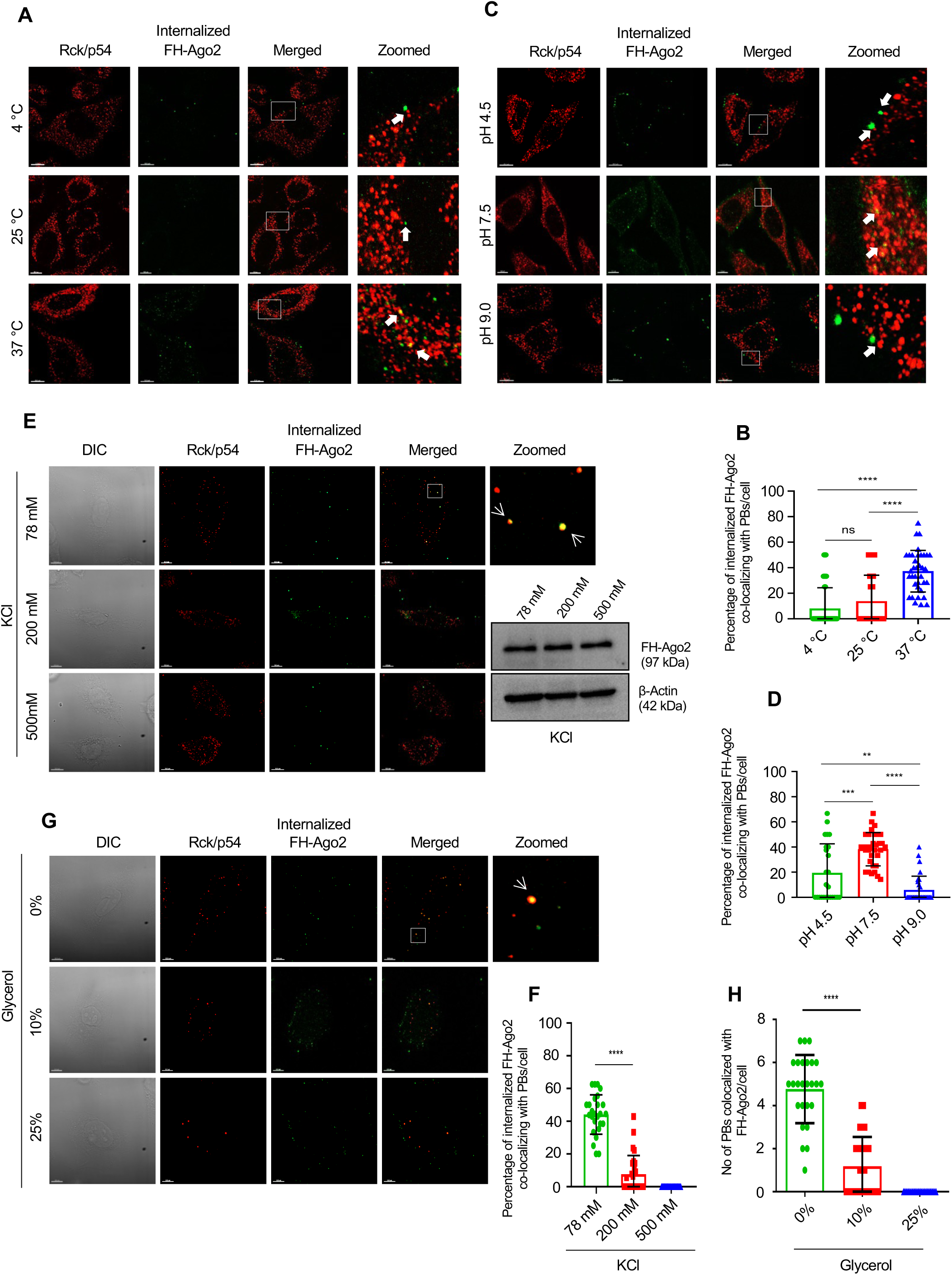
Phase separation and P-body targeting of Ago2 is dependent on osmolarity and viscosity of the incubation buffer. **(A)** Confocal Images showing co-localization between internalized FH-Ago2 protein (Green) and endogeneousRck/p54 protein (Red) in HeLa cells (recipient) incubated with FH-Ago2 transfected HEK293 cell (donor) lysate at 4 °C, 25 °C, 37 °C. **(B)** Percentageof internalized FH-Ago2 protein (Green) co-localizing withendogenous Rck/p54 protein (Red) in HeLa cell (recipient) incubated with FH-Ago2 transfected HEK293 cell (donor) lysate at 4 °C, 25 °C, 37 °C. (p=0.2439(between 4 °C and 25 °C groups), p <0.0001 (between 25 °C and 37 °C groups), p < 0.0001 (between 4 °C and 37 °C groups), n=3, >25 cells used for quantification). **(C)** Confocal Images showing co-localization between internalized FH-Ago2 protein (Green) and endogenous Rck/p54 protein (Red) in HeLa cells (recipient) incubated with FH-Ago2 transfected HEK293 cell (donor) lysate in lysis buffer having pH4.5, 7.5, 9.0. **(D)**Percentage of internalized FH-Ago2 protein (Green) co-localizing with endogenous Rck/p54 protein (Red) in HeLa cell (recipient) incubated with FH-Ago2 transfected HEK293 cell (donor) lysate, lysed with lysis buffer having pH4.5, 7.5, 9.0 (p=0.0002(between pH=4.5 and pH=7.5 groups), p < 0.0001 (between pH=7.5 and pH=9 groups), p=0.0054(between pH=4.5 and pH=9 groups), n=3, >25cells used for quantification). **(E)** Confocal Images showing co-localization between internalized FH-Ago2 protein (Green) and endogenous Rck/p54 protein (Red) in HeLa cells (recipient) incubated with FH-Ago2 transfected HEK293 cell (donor) lysates containing 78, 200, or 500 mM KCl. Western blot of FH-Ago2 detected by anti-HA antibody in lysate after incubation with different salt concentration. β-actin used as loading control. **(F)**Percentage of internalized FH-Ago2 protein (Green) co-localizing with endogenous Rck/p54 protein (Red) in HeLa cell (recipient) incubated with FH-Ago2 transfected HEK293 cell (donor) lysates containing 78, 200, 500 mM KCl (p < 0.0001 (between 78 mM and 200 mM groups), n=3, >25 cells used for quantification). **(G)** Confocal Images showing co-localization between internalized FH-Ago2 protein (Green) and Rck/p54 protein (Red) in HeLa cells (recipient) incubated with FH-Ago2 transfected HEK293 cell (donor) lysates containing 0, 10, or 25 % Glycerol. **(H)** Quantification of co-localization between internalized FH-Ago2 protein (Green) and Rck/p54 protein (Red) in HeLa cell (recipient) incubated with FH-Ago2 transfected HEK293 cell (donor) lysates with 0, 10, 25 % Glycerol (p < 0.0001 (between 0% and 10% groups), n=3, >25 cells used for quantification). Merged images and images obtained by differential interference contrast (DIC) are also shown. Zoomed images are magnified representation of the inset box in merged images. Fields have been detected in 63X magnification. Scale bar represents 10 µm. Data represents means ± SDs; ns, non-significant, *p < 0.05, **p < 0.01, ***p < 0.001, ****p < 0.0001. p values were obtained by using two-tailed unpaired Student’s t test.

### ATP hydrolysis is required for Ago2 phase separation and PB-localization

What is the energy requirement of PB-targeting and Ago2 body formation? We hypothesized ATP, which acts as energy source in many biological processes, may also have a role in regulation of protein solubility and compartmentalization (19). Thus, ATP should impact Ago2 phase separation process. We treated the donor HEK293 cell lysate for 1 hour at 30 °C with 0.5 U/ml of Apyrase enzyme to hydrolyze ATP present in the lysate. We incubated the ATP-depleted lysate with the recipient HeLa cells and observed, a decreased Ago2 internalization and PB targeting in ATP-depleted condition. Interestingly, supplementation of Apyrase treated lysate with ATP but not with AMP-PNP (an non-hydrolysable analogue of ATP), increased the FH-Ago2 targeting to PBs (Figure 4A-C). If we treat the donor HEK293 cell with 3 µM FCCP or 1 µM Oligomycin for 1 hour at 37 °C before lysis to reduce the ATP level of the donor cells by decreasing oxidative phosphorylation, we documented a reduction in PB targeting of FH-Ago2 upon incubation with lysate with a decreasing concentration of ATP due to FCCP or oligomycin treatment of donor cells. Therefore, compared to control condition, we did observe a significant decrease in the PB phase separation of FH-Ago2 on ATP depletion or reduction in its concentration. The ATP depletion by FCCP or oligomycin treatment did not affect the stability and content of FH-Ago2 present in the lysate after incubation (Figure 4D). To confirm that the ATP depletion could affect dynamics of PBs and thus can alter Ago2 compartmentalization, we did a FRAP experiment where we recorded a time lapse of fluorescence recovery after photo bleaching of exogenously expressed GFP-Dcp1a bodies in recipient HeLa cells after 30 mins of incubation with donor HEK293 cell lysate treated with Apyrase. We observed a reduced recovery of fluorescence of GFP-Dcp1a bodies post photo bleaching in Apyrase treated condition as compared to the control set (Figure 4G and 4H). How the ATP hydrolysis is used for PB-targeting of Ago2? It was reported before that HSP90 may have a role to play in Ago2 activity regulation as it is known to interact with Ago2 protein (20). With HSP90 inhibition in the donor HEK293 cell lysate, by using Geldanamycin during the incubation, we did not find any effect of HSP90 inhibition on Ago2 localization to PBs (Supplementary Figure S2B). The other possible effect of ATP depletion on PB targeting of Ago2 may be caused by altered intracellular trafficking of Ago2 happening due to impaired cytoskeleton dynamics in ATP reduced state. We explored the effect of drugs that depolymerizes microtubule structures and thus should alter the intracellular Ago2 movement like it happening in ATP-reduced condition. To test this, the recipient HeLa cells were treated with 1.5 or 3 µM of Nocodazole for 2 hours at 37°C before incubation with donor HEK293 cell lysate. This treatment reduced the internalization as well as transport of the Ago2 proteins to the endogenous PBs (Figure 4E and 4F). Although Ago2 targeting to PBs gets altered with changes in viscosity and osmolarity of the incubation buffer, we did not find a change in the levels of soluble FH-Ago2 present in the lysate after high salt, different pH or viscosity to justify the differences observed in Ago2 localization to PBs with altered reaction conditions (Figure 3E and unpublished data). We used H_2_O_2_ to induce oxidative stress in HeLa cells, but we did not find any effect of stress induction on internalized Ago2 compartmentalization to PBs (Supplementary Figure S2C). For LLPS, another important parameter that could have most profound effect is the concentration of protein used in the assay. After treatment with increasing concentration of Proteinase K for 10 mins at 37 °C, the donor HEK293 cell lysate formed a smaller number of internalized Ago2 bodies with reduced a co-localization with PBs which were also getting significantly lowered in numbers with increasing concentration of Proteinase K treatment (Supplementary Figure S3A, S3D, S3E and S3F). This data suggests that the phase separated Ago2 are sensitive to Proteinase K. Therefore PB targeting is possibly reversible and unlike a prion-like transition, Ago2 remains protease sensitive after the phase separation to PBs (21). Compared to internalized Ago2 pool, the Proteinase K mediated degradation of FH-Ago2 present in the lysate suggests that the soluble protein in the lysate is similarly sensitive to proteinase K treatment compared to PB-phase separated FH-Ago2 (Supplementary Figure S3B and S3C).

**Figure 4.**
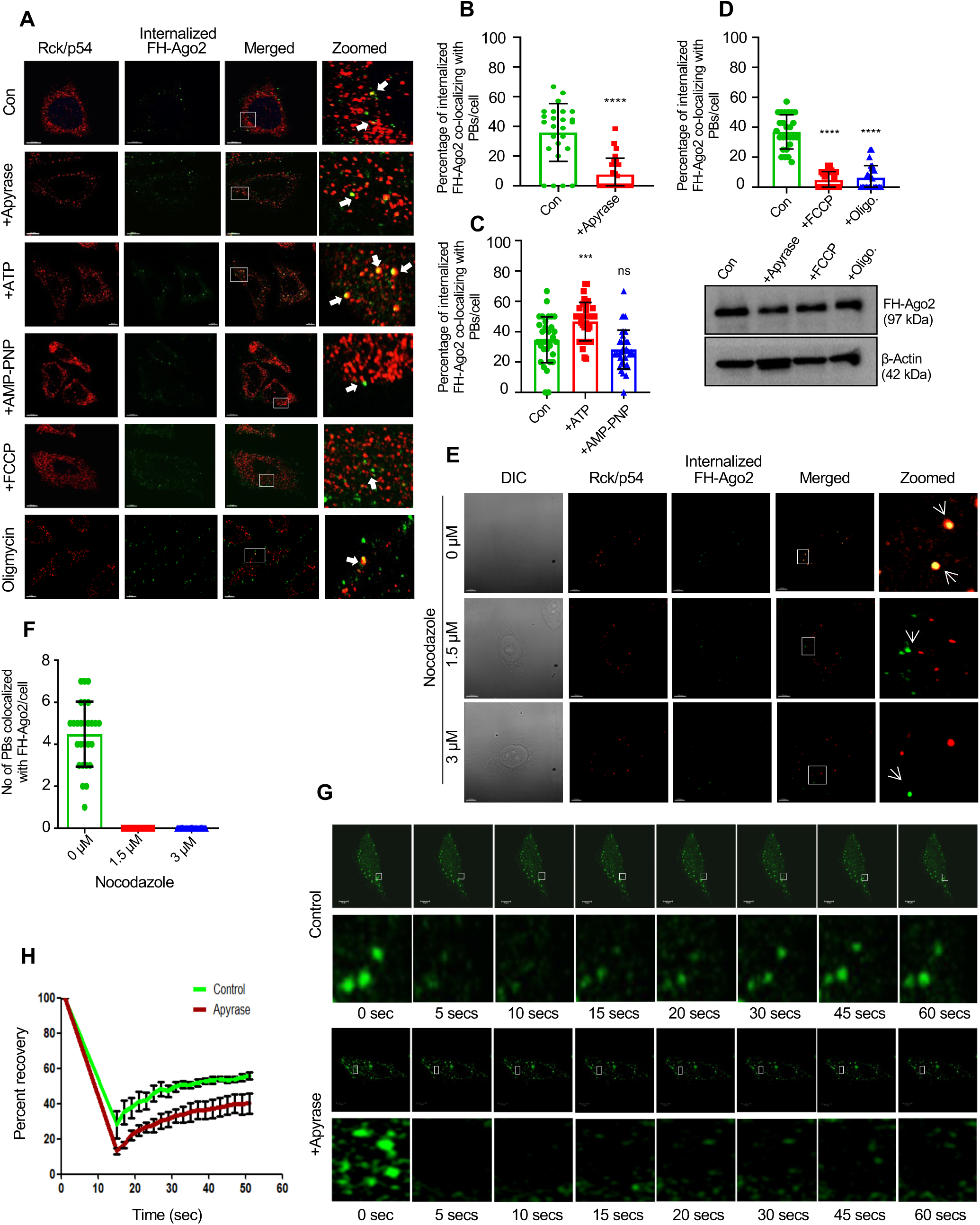
Phase separation and P-body targeting of Ago2 requires ATP hydrolysis. **(A)** Confocal Images showing co-localization between internalized FH-Ago2 protein (Green) and endogenous Rck/p54 protein (Red) in HeLa cells (recipient) incubated with FH-Ago2 transfected HEK293 cell (donor) control (untreated) lysate, or lysate treated with 0.5 U/ml Apyrase enzyme for 1 hour at 30 °C, or donor cell lysate supplemented with 1 mM of ATP, or AMP-PNP, or lysates prepared from HEK293 donor cells treated with 3 µM FCCP or 1 µM Oligomycin at 37 °C for 1 hour before cell lysis. **(B-D)**Percentage of internalized FH-Ago2 protein (Green) co-localizing with endogenous Rck/p54 protein (Red) in HeLa cell (recipient) incubated with FH-Ago2 transfected HEK293 cell (donor) lysate in the abovementioned conditions. (Between control and Apyrase treated groups)p < 0.0001, n=3, >25 cells used for quantification (**B**), p=0.0007 (between control and ATP supplemented groups), p=0.0594 (between control and AMP-PNP supplemented groups), n=3, >30cells used for quantification (**C**), p < 0.0001 (between control and FCCP treated groups), p < 0.0001 (between control and Oligomycin treated groups), n=3, >25 cells used for quantification (**D**). Western blot of FH-Ago2 was done with anti HA antibody to measure level of FH-Ago2 after the treatment of lysate with Apyrase, FCCP or oligomycin. β-actin used as loading control.**(E)** Confocal Images showing co-localization between internalized FH-Ago2 protein (Green) and endogenousRck/p54 protein (Red) in HeLa cells (recipient), priorly treated with 0, 1.5, 3 µM Nocodazole for 2 hours at 37 °C, followed by incubation with FH-Ago2 transfected HEK293 cell (donor) lysate. **(F)** Quantification of co-localization between internalized FH-Ago2 protein (Green) and endogenous GFP-Dcp1a protein (Red) in HeLa cell (recipient), priorly treated with Nocodazole and incubated with FH-Ago2 transfected HEK293 cell (donor) lysate. **(G)** Timelapse images of recipient HeLa cells, transiently transfected with GFP-Dcp1a, captured during FRAP analysis after 30 mins of incubation with HEK293 (donor) cell lysate in control and 0.05 U/ml Apyrase treated condition (for 1 hour at 30 °C). Recovery was captured till 60 seconds. **(H)** Percent recovery of fluorescence after photobleaching of GFP-Dcp1a bodies in control and 0.05 U/ml Apyrase treated condition (for 1 hour at 30 °C). Merged images and images obtained by differential interference contrast (DIC) are given. Zoomed images are magnified representation of the inset box in merged images. Fields have been detected in 63X magnification. Scale bar represents 10 µm. Data represents means ± SDs; ns, non-significant, *p < 0.05, **p < 0.01, ***p < 0.001, ****p < 0.0001. p values were obtained by using two-tailed unpaired Student’s t test.

### *In vitro* PB-targeting of mRNA depends on miRNA and Ago2 binding

Do miRNA targeted mRNAs are also getting compartmentalized to PBs as observed in the *in vivo* context in mammalian cells? Do they accompany Ago2 in PB-compartmentalization? We observed compartmentalization of internalized miRNAs to Ago2 bodies in the recipient HeLa cell expressing GFP-Ago2 after incubation with donor HEK293 cell lysate supplemented with the synthetic sense strand of Cy3-tagged miR-122. We performed the assay with increasing concentration of miR-122 and found appreciable level of miR-122 compartmentalized at 20 nM concentration. Targeting of internalized miR-122 to GFP-Dcp1a bodies (another P-body marker protein) was also observed at the same concentration of Cy3-tagged miR-122 (Figure 5A and 5B). Next, we intended to see whether PB compartmentalization of miRNA-targeted mRNAs also happens in vitro in this digitonin based cell permeabilization assay. To explore the importance of miRNA associated proteins like Ago2 in this process, we had taken *in vitro* transcribed m7G-capped and polyadenylated RL-3xB-let-7a reporter mRNA having three bulged let-7a binding sites. We supplemented the donor HEK293 cell lysate with 100 nM of this mRNA and incubated with the GFP-Dcp1a expressing recipient HeLa cell, followed by RNA-FISH using Cy3-labeled oligos against RL sequence (22)(Figure 5C). We performed this assay with lysate of HEK293 in HeLa cells expressing pCIneo, NHA-LacZ (act as control), NHA-GW182 or FH-Ago2. While no significant difference was observed upon NHA-GW182 expression as compared to the control, the let-7a repressed target reporter mRNAs localization to PBs appeared to be enhanced by FH-Ago2 expression (Figure 5D, 5E and 5F).

**Figure 5.**
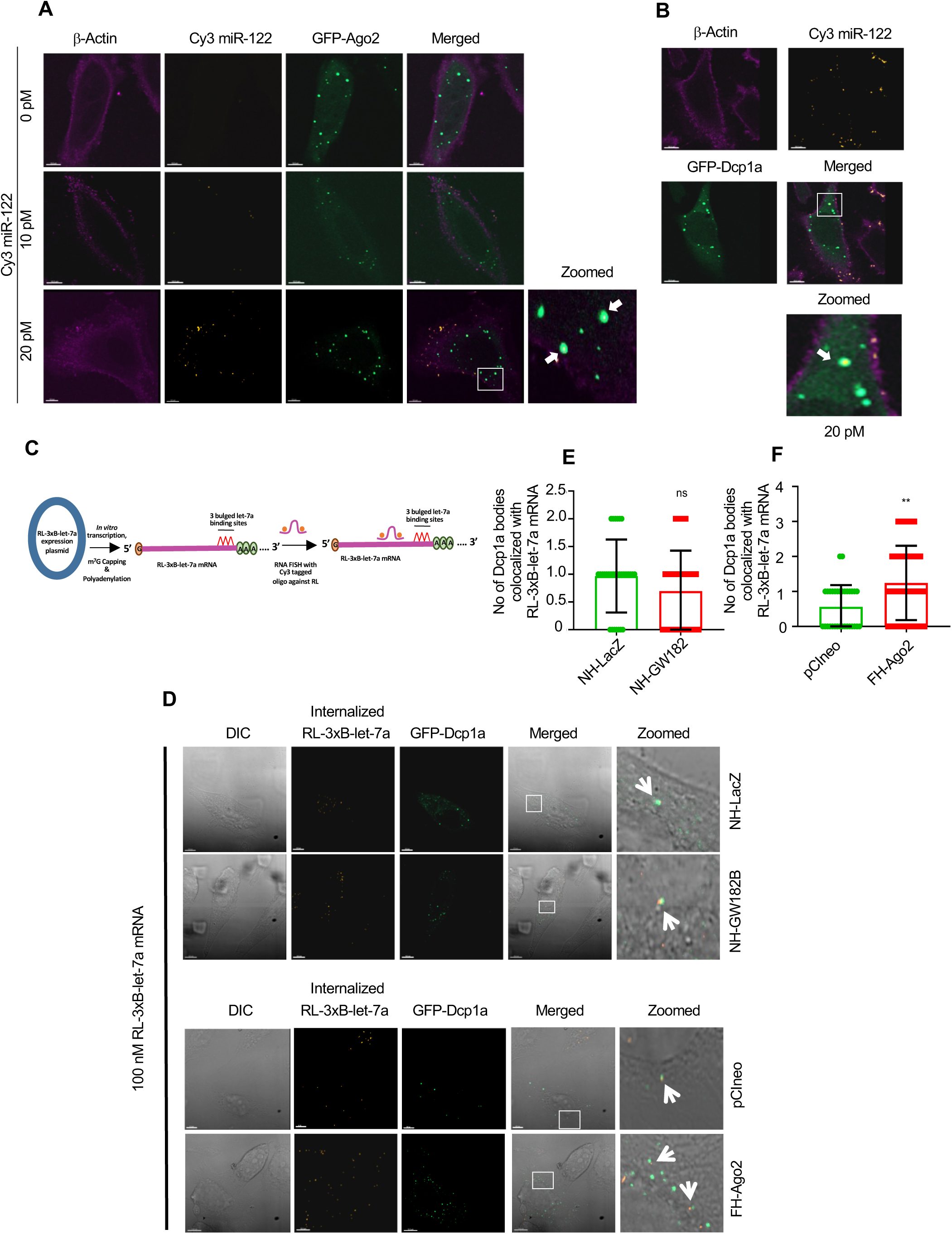
miRNA and mRNAs are targeted to PBs with Ago2 *in vitro*. **(A)** Confocal Images showing co-localization between internalized synthetic Cy3-miR122 sense strand (orange) and transfected GFP-Ago2 protein (Green) in HeLa cells (recipient) incubated with transfected HEK293 cell (donor) lysate supplemented with 0, 10, 20 pM of the synthetic miRNA. **(B)** Confocal Images showing co-localization between internalized synthetic Cy3-miR122 sense strand (orange) and exogenously transfected GFP-Dcp1a protein (Green) in HeLa cells (recipient) incubated with untransfected HEK293 cell (donor) lysate supplemented with 20 pM of the synthetic miRNA. **(C)** A schematic representation of strategy of in situ detection of RL-3xB-let-7a IVT RNA in recipient cells after the cell-permeabilization-resealing assay followed by RNA-FISH. **(D)** Confocal Images showing co-localization between internalized RL-3xB-let-7a mRNA visualized by RNA-FISH (orange) and exogenously expressed GFP-Dcp1a protein (Green) in HeLa cells (recipient) co-transfected with NH-LacZ or NH-GW182 and pCI-neo or FH-Ago2 expression plasmids (NH-LacZ and pCI-neo taken as control plasmids) and incubated with untransfected HEK293 cell (donor) lysate supplemented with 100 nM of RL-3xB-let-7a mRNA. **(E-F)** Quantification of the co-localization between internalized RL-3xB-let-7a mRNA (Orange) and exogenously expressed GFP-Dcp1a protein (Green) in HeLa cell (recipient) co-transfected with NH-LacZ or NH-GW182 expression plasmids (p=0.1243, n=3, >30 cells used for quantification, E) and pCI-neo or FH-Ago2 expression plasmids (p=0.0026, n=3, >30 cells used for quantification, F). Merged images and images obtained by differential interference contrast (DIC) are given. Zoomed images are magnified representation of the inset box in merged images. Fields have been detected in 63X magnification. Scale bar represents 10 µm. Data represents means ± SDs; ns, non-significant, *p < 0.05, **p < 0.01, ***p < 0.001, ****p < 0.0001. p values were obtained by using two-tailed unpaired Student’s t test.

Interestingly, presence of RNA also appeared to be crucial for the phase separation of Ago2 to PBs. Upon RNaseA treatment, that degrades RNA in lysates, the localization of Ago2 to PBs was greatly affected. Localization of internalized Ago2 proteins to endogenous PBs was significantly retarded when the recipient HeLa cell was incubated with FH-Ago2 transfected donor HEK293 cell lysate treated with increasing concentration of RNase A (Supplementary Figure S4A and S4B). However, when the lysate was treated with DNase I, no effect on PB localization of Ago2 was observed (Supplementary Figure S4C and S4D). We went on to supplement FH-Ago2 expressing HEK293 lysate with RL-Con or RL-3xB-let-7a mRNAs, however, no appreciable changes in the internalized Ago2 localization to the GFP-Dcp1a bodies were observed (Supplementary Figure S5A and C). However, translation inhibition by Puromycin that generates a pool of run-off mRNAs resulted in an enhanced PB size *in vitro*. Cycloheximide, that arrests translating polysomes had an opposite effect on the same. Therefore, this data along with previous findings (5) suggests that Ago2 targeting or Phase separation get enhanced when the Ago2 bound mRNAs are not part of translational machinery and miRNA binding may favor the equilibrium to form ribosome free mRNA-Ago2 complex to drive the mRNA and Ago2 entry to PBs *in vitro* (Figure S5B, S5D and S5E). Interestingly the run-off mRNA availabilityby puromycin treatment doesn’t increase the number of PBs and therefore the mRNA may get targeted to bodies that are already positive for Ago2 miRNPs and non-polysomal mRNA alone can’t initiate de novo body formation. Thus, the mRNA targeting to PBs happens possibly due to Ago2-miRNP binding of the mRNAs.

To further explore the role of miRNA on target message compartmentalization to PBs, we prepared *in vitro* transcribed RL-3xB-miR-122 mRNA and supplemented it to HEK293 lysate which was incubated with GFP-Dcp1a expressing recipient HeLa cells co-transfected with pmiR-122 plasmid to express this liver specific miRNA in Hela (Figure 6A). Following RNA-FISH technique, we had observed PB compartmentalization of RL-3xB-miR-122 mRNA only in cells expressing miR-122 which was not detected in PBs in absence of miR-122 expression (Figure 6B and C). Therefore, translationally inhibited, and miRNA-targeted mRNAs may have a positive influence on PB targeting of Ago2 and the mRNAs themselves also are getting trafficked to PBs when repressed by miRNAs. Therefore, miRNAs are responsible for target mRNA compartmentalization to PBs. Does miRNA binding is essential for Ago2 protein for its compartmentalization to PBs?

**Figure 6.**
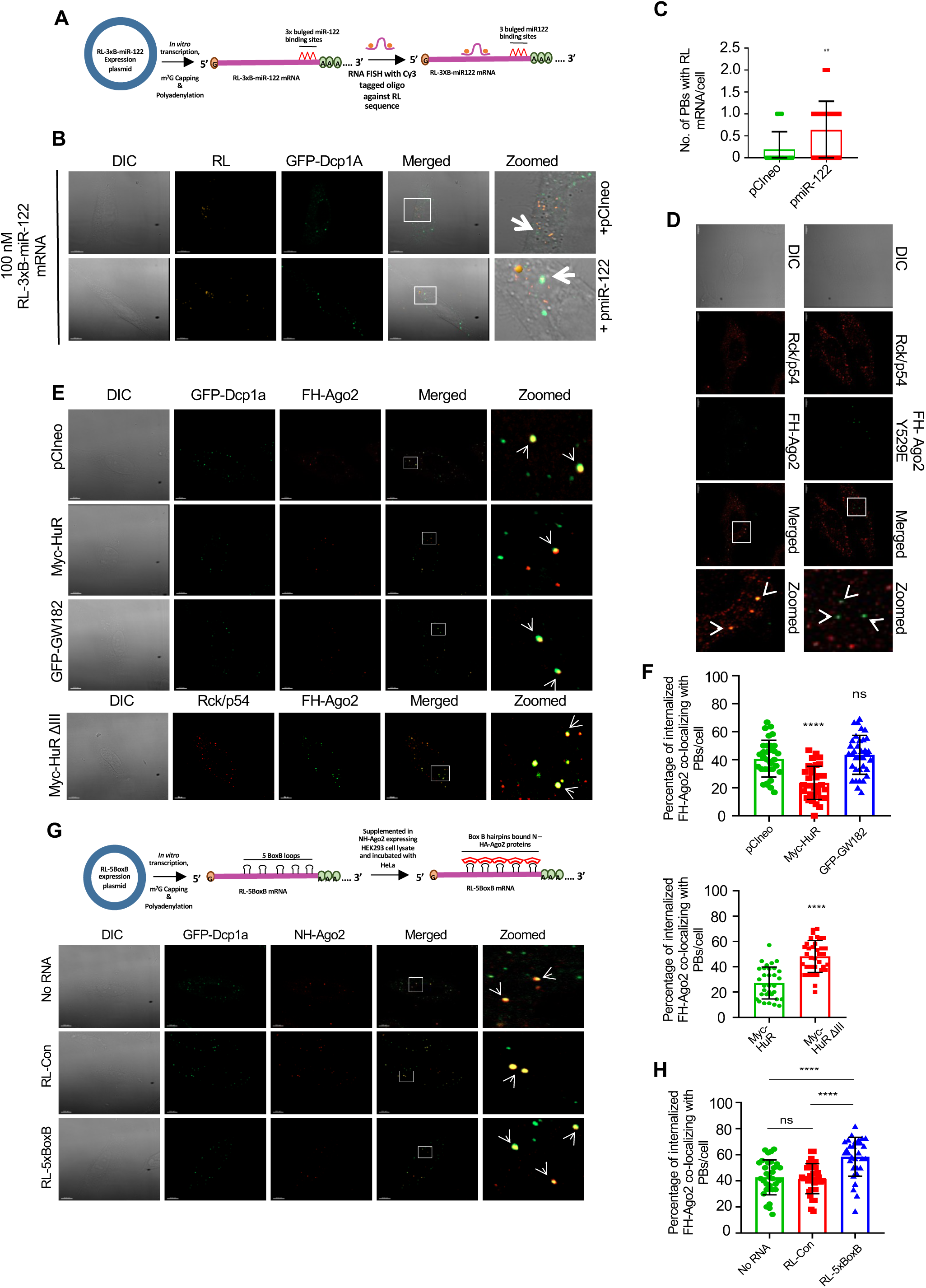
Phase separation and P-body targeting of Ago2 is dependent miRNA binding of Ago2. **(A)** A schematic representation of RL-3xB-miR-122 RNA preparation and its use in the cell-permeabilization-resealing assay followed by RNA-FISH for detection. **(B)** Confocal Images showing co-localization between internalized RL-3xB-miR-122 mRNA visualized by RNA-FISH (orange) and exogenously expressed GFP-Dcp1a protein (Green) in HeLa cells (recipient) expressing or not expressing pmiR-122 and incubated with HEK293 cell (donor) lysate supplemented with 100 nM of RL-3xB-miR-122 mRNA. **(C)** Quantification of co-localization between internalized RL-3XB-miR-122 mRNA (Orange) and exogenously expressed GFP-Dcp1a protein (Green) in control and pmiR-122 expressing HeLa cell (recipient) incubated with untransfected HEK293 cell (donor) lysate supplemented with 100 nM of RL-3xB-miR-122 mRNA (p=0.0019, n=3, >30 cells used for quantification). **(D)**Confocal images showing co-localization between internalized FH-Ago2 protein or FH-Ago2 Y529E mutant protein (Green) and endogenously immunostainedRck/p54 protein (Red) in HeLa cells (recipient) incubated with FH-Ago2 or FH-Ago2 Y529E transfected HEK293 cell (donor) lysate. **(E)** Confocal images showing co-localization between internalized FH-Ago2 protein (Red) and exogenously expressed GFP-Dcp1a protein (Green) in HeLa cells (recipient) incubated with FH-Ago2 transfected HEK293 cell (donor), co-transfected with either pCIneo or Myc-HuR or Myc-HuRΔIII orGFP-GW182, lysate. **(F)**Percentage of internalized FH-Ago2 protein (Red) co-localizing with exogenously expressed GFP-Dcp1a protein (Green) in HeLa cell (recipient) incubated with FH-Ago2 transfected HEK293 cell (donor), co-transfected with either pCIneo or Myc-HuR or Myc-HuRΔIII orGFP-GW182, lysate (p < 0.0001, (between pCIneo and Myc-HuR groups), p=0.4072(between pCIneo and GFP-GW182 groups), n=3, >30 cells used for quantification). In the bottom panel relative levels of PB-localized FH-Ago2 in presence of Myc-HuR and Myc-HuRΔIII have been depicted(p < 0.0001, n=3, >30 cells used for quantification). **(G)** Confocal images showing co-localization between internalized NH-Ago2 protein (Red) and exogenously expressed GFP-Dcp1a protein (Green) in HeLa cells (recipient) incubated with NH-Ago2 transfected HEK293 cell (donor) lysate supplemented with no RNA or 100 nM of RL-con or RL-5BoxB mRNA. **(H)**Percentage of internalized FH-Ago2 protein (Red) co-localizing with exogenously expressed GFP-Dcp1a protein (Green) in HeLa cells (recipient) incubated with NH-Ago2 transfected HEK293 cell (donor) lysate supplemented with no RNA or 100 nM of RL-con or RL-5BoxB mRNA (p<0.0001(between no RNA and RL-5BoxB mRNA groups), p<0.0001 (between RL-con and RL-5BoxB mRNAs groups), p=0.7434 (between no RNA and RL-con mRNA groups), n=3, >30 cells used for quantification). Merged images and images obtained by differential interference contrast (DIC) are given. Zoomed images are magnified representation of the inset box in merged images. Fields have been detected in 63X magnification. Scale bar represents 10 µm. Data represents means ± SDs; ns, non-significant, *p < 0.05, **p < 0.01, ***p < 0.001, ****p < 0.0001. p values were obtained by using two-tailed unpaired Student’s t test.

### miRNA binding of Ago2 is necessary for phase separation to PBs and is inhibited by miRNA “sponge” protein HuR

To explore the requirement of miRNA binding of Ago2 for its phase separation to PBs, we tested the compartmentalization of wild type and mutant Ago2 that could not bind miRNA. Phase separation of internalized Ago2 proteins seems to be dependent on its association with miRNA, as we have observed a retarded targeting of a mutant Ago2 protein FH-Ago2Y529E having a tyrosine to glutamic acid substitution mutation at 529 position which removes the ability of Ago2 to bind miRNA to form miRISC complex(23). This mutation reduces FH-Ago2Y529E targeting to P-bodies when we incubated the recipient HeLa cell with the wild type and mutant FH-Ago2 protein expressing donor HEK293 cell lysate (Figure 6D). HuR, an RNA binding protein belonging to ELAV family, has been reported previously to be involved in derepression of miRNA-target messages by binding to the AU-rich sequence on the 3’ UTR of the target mRNAs and thereby replaces the miRNPs from cognate mRNAs (10). miRNAs also can bind with HuR. The protein HuR, by binding with the miRNAs, removes them from Ago2-miRNA complex to derepress the target mRNA and also accelerates miRNA export via extracellular vesicles (24). On contrary, GW182 protein promotes miRNA stability possibly by affecting its compartmentalization to PBs in mammalian cells (25). We wanted to find out the effect of HuR or GW182 protein expression on the P-body targeting of internalized Ago2 proteins. For this, we had co-transfected the GFP-Dcp1a expressing recipient HeLa cells with pCIneo, Myc-HuR or GFP-GW182 expression plasmids and incubated with FH-Ago2 expressing donor HEK293 cell lysate. We observed, in Myc-HuR expressing HeLa cells, Ago2 targeting to PBs got significantly decreased, while no effect was seen upon GFP-GW182 expression (Figure 6E and 6F). As HuR binds and removes miRNA from Ago2, this data suggests that the miRNA binding of Ago2 is essential for PB phase separation. However, the PB targeting of Ago2-miRNA complex may get accelerated upon translational repression of the target mRNAs. We performed P-body targeting assay with*in vitro* transcribed mRNA RL-5BoxB having 5BoxB RNA element and used it to supplement the NH-Ago2 expressing donor HEK293 cell lysate for incubation with the GFP-Dcp1a expressing recipient HeLa cell. This RL-5BoxB RNA can bind to the N protein of NH-tagged Ago2 protein. We observed an increased Ago2 compartmentalization to P-bodies of NH-Ago2 in presence of RL-5BoxB mRNA compared to what observed in no exogenous RNA or RL-con mRNA supplementation (Figure 6G and 6H). This data suggests that mRNA binding of Ago2, either by miRNA or by direct tethering of the NH-Ago2 to an mRNA, accelerates PB targeting of Ago2. Therefore, the phase transition of Ago2 is notably getting affected by several factors including the presence of target mRNAs not associated with translating polysomes.

### Amyloidogenic proteins accelerate PB compartmentalization of Ago2 miRNPs

We have documented how the phase separation of Ago2 protein is affected by several physiological parameters. We wanted to check how the presence of amyloidogenic proteins could affect Ago2 compartmentalization to PBs. It has been documented before that amyloid beta (Aβ) 1-42 amino acid long oligomers affect miRNA activity in glial cells and that was accompanied by an increased PB numbers (17). The depletion of PBs by using siRNAs against PB components affects the miRNA activity and suppresses expression of inflammatory cytokines in Aβ_1-42_treated glial cells. This data suggests PB localization of miRNPs may be important for Ab_1-42_-mediated miRNA activity modulation and that may happen due to increased phase separation of Ago2 to PBs in presence of amyloidogenic proteins. We have used Aβ_1-42_ at increasing concentration to see its effect on PB localization of FH-Ago2 in HEK293 lysate treated permeabilized HeLa cells (Figure 7A). We documented increased Ago2 bodies and PB-localization of Ago2 in presence of Aβ_1-42_ at 2.5 µm concentration. However non-amyloidogenic Aβ_(1-40)_ does not show any effect on PB localization of Ago2. Both Aβ_(1-42)_ and Aβ_(1-40)_ did not have any effect on existing PB number in this *in vitro* assay (Figure 7B-E). We have observed the similar result with non-permeabilized HeLa cells treated with similar concentrations of Aβ_1-42_ for 24 hours (Figure S6A and B). When we immunostained internalized Aβ_1-42_ in recipient GFP-Dcp1a expressing HeLa cells incubated with FH-Ago2 transfected donor HEK293 cell lysate along with 2.5 µm Aβ_(1-42),_ we have seen that internalized Aβ_(1-42)_ localize to PBs and increase PBs co-localization with internalized FH-Ago2 (Figure S6C and D). The Myc-HuR expression that can remove miRNA from Ago2, caused a decreased localization of FH-Ago2 to PBs in presence of Aβ_1-42_ oligos (Figure 7F). Interestingly, cy3-miR-122 localization to PBs get enhanced in presence of Aβ_(1-42)_ oligomers (Figure 7G and H).

**Figure 7.**
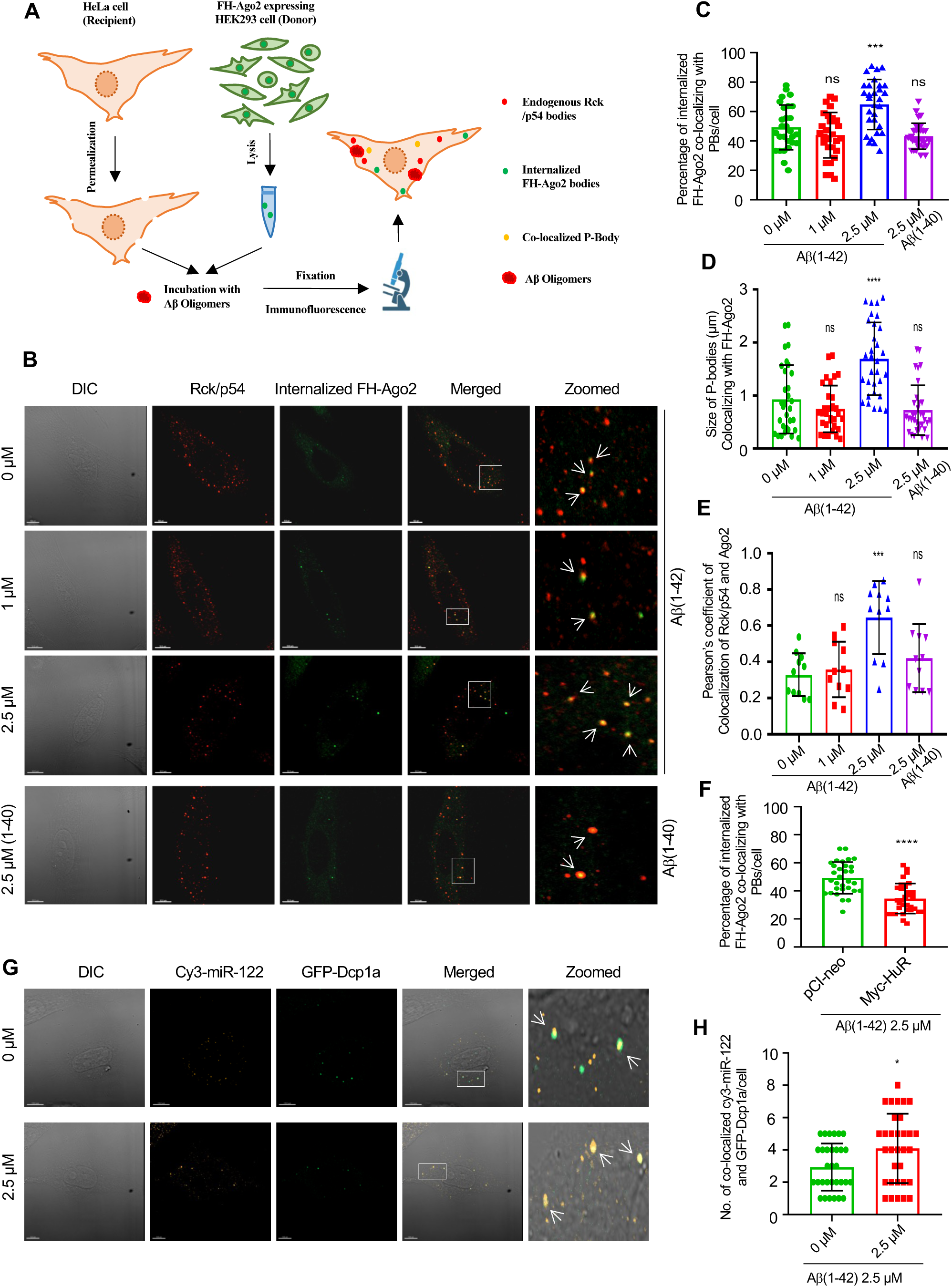
Amyloid beta oligomers facilitate compartmentalization of Ago2 to RNA processing bodies. **(A)** A schematic representation of the cell permeabilization-resealing assay using HeLa cells as recipient and HEK293 cells as donor including Aβ treatment. (**B**) Confocal Images showing co-localization between internalized FH-Ago2 protein (Green) and endogenous Rck/p54 protein (Red) in HeLa cells (recipient) incubated with FH-Ago2 transfected HEK293 cell (donor) lysate along with 0, 1, 2.5 µM of Aβ_(1-42)_ and 2.5 µM of Aβ_(1-40)_. Here, Aβ_(1-40)_ used as control that does not show pathogenicity. **(C)** Percentage of internalized FH-Ago2 protein (Red) co-localizing with endogenous Rck/p54 protein (Red) in HeLa cell (recipient) incubated with FH-Ago2 transfected HEK293 cell (donor) lysate treated with 0, 1, 2.5 µM of Aβ_(1-42)_ and 2.5 µM of Aβ_(1-40)_. (p=0.1964 (between 0 µM and 1 µM Aβ_(1-42)_ groups), p=0.0006 (between 0 µM and 2.5 µM Aβ_(1-42)_ groups), p=0.0644 (between 0 µM and 2.5 µM Aβ_(1-40)_ groups), n=3, >25cells used for quantification). **(D)** Size of co-localized internalized FH-Ago2 protein (Green) and endogenous Rck/p54 protein (Red) in HeLa cell (recipient) incubated with FH-Ago2 transfected HEK293 cell (donor) lysate treated with 0, 1, 2.5 µM of Aβ_(1-42)_ and 2.5 µM of Aβ_(1-40)._ (p=0.2026 (between 0 µM and 1 µM Aβ_(1-42)_ groups), p < 0.0001 (between 0 µM and 2.5 µM Aβ_(1-42)_ groups), p=0.1524 (between 0 µM and 2.5 µM Aβ_(1-40)_ groups), n=3, >30 cells used for quantification). **(E)** Pearson’s coefficient of colocalization between internalized FH-Ago2 protein (Green) and endogenous Rck/p54 protein (Red) in HeLa cell (recipient) incubated with FH-Ago2 transfected HEK293 cell (donor) lysate treated with 0, 1, 2.5 µM of Aβ_(1-42)_ and 2.5 µM of Aβ_(1-40)_. (p=0.6229 (between 0 µM and 1 µM Aβ_(1-42)_ groups), p=0.0002 (between 0 µM and 2.5 µM Aβ_(1-42)_ groups), p=0.1861 (between 0 µM and 2.5 µM Aβ_(1-40)_ groups), n=3). (**F**) Percentage of internalized FH-Ago2 protein (Red) co-localizing with endogenous Rck/p54 protein (Red) in HeLa cell (recipient) incubated with FH-Ago2 transfected HEK293 cell (donor), co-transfected with pCI-neo or myc-HuR, lysate along with 2.5 µM of Aβ_(1-42)_ (p<0.0001, n=3, >30 cells used for quantification). (**G**) Confocal Images showing co-localization between internalized synthetic sense strand of cy3-miR-122 (orange) and exogenously expressed GFP-Dcp1a protein (Green) in HeLa cells (recipient) incubated with untransfected HEK293 cell (donor) lysate supplemented with 20 pM of synthetic sense strand of cy3 tagged miR-122 and treated with 0 or 2.5 µM of Aβ_(1-42)_. (**H**) Quantification of co-localization between internalized synthetic sense strand of cy3-miR-122 (orange) and exogenously expressed GFP-Dcp1a protein (Green) in HeLa cells (recipient) incubated with untransfected HEK293 cell (donor) lysate supplemented with 20 pM of synthetic sense strand of cy3 tagged miR-122 and treated with 0 or 2.5 µM of Aβ_(1-42)_. (p=0.0152, n=3, >30 cells used for quantification). Merged images and images obtained by differential interference contrast (DIC) are given. Zoomed images are magnified representation of the inset box in merged images. Fields have been detected in 63X magnification. Scale bar represents 10 µm. Data represents means ± SDs; ns, non-significant, *p < 0.05, **p < 0.01, ***p < 0.001, ****p < 0.0001. p values were obtained by using two-tailed unpaired Student’s t test.

### Intracellular dynamics and diffusion of phase separated Ago2 bodies get decreased on Amyloid beta oligomers treatment which get rescued in presence of Myc-HuR

To determine the intracellular dynamics of GFP-Ago2 in HeLa cell we adopted Fluorescence Correlation spectroscopy (FCS) measurement to follow the temporal fluctuation of fluorescence intensity inside the small confocal volume (∼ femtolitre). Exploration of the autocorrelation function obtained from FCS measurement provides us the diffusion coefficient (D) of the molecule by using a suitable model. Ago2 protein was tagged with fluorophore GFP. To explore GFP-Ago2 dynamics inside the puncta formed in HeLa cell we used region specific point-FCS (Figure 8A and B). At first, we measured diffusion coefficient of GFP-Ago2 both in the absence and presence of Aβ_(1-42)_. GFP-Ago2 expressing Hela cells (recipient) were permeabilized and incubated with untransfected HEK293 (donor) cell lysate in the absence or presence of 2.5 µM Aβ_(1-42)_. The shift in auto-correlation function of GFP-Ago2 in presence of Aβ_(1-42)_ indicates an arrested dynamics of GFP-Ago2 inside the cell. (Figure 8C and D). The mean diffusion coefficient of GFP-Ago2 in the presence of Aβ_(1-42)_ inside the puncta was found to be ∼ 2 times lower compared to only GFP-Ago2 puncta, which further suggests mobility of GFP-Ago2 is less in the presence of Aβ_(1-42)_. (Figure 8C and D)

**Figure 8.**
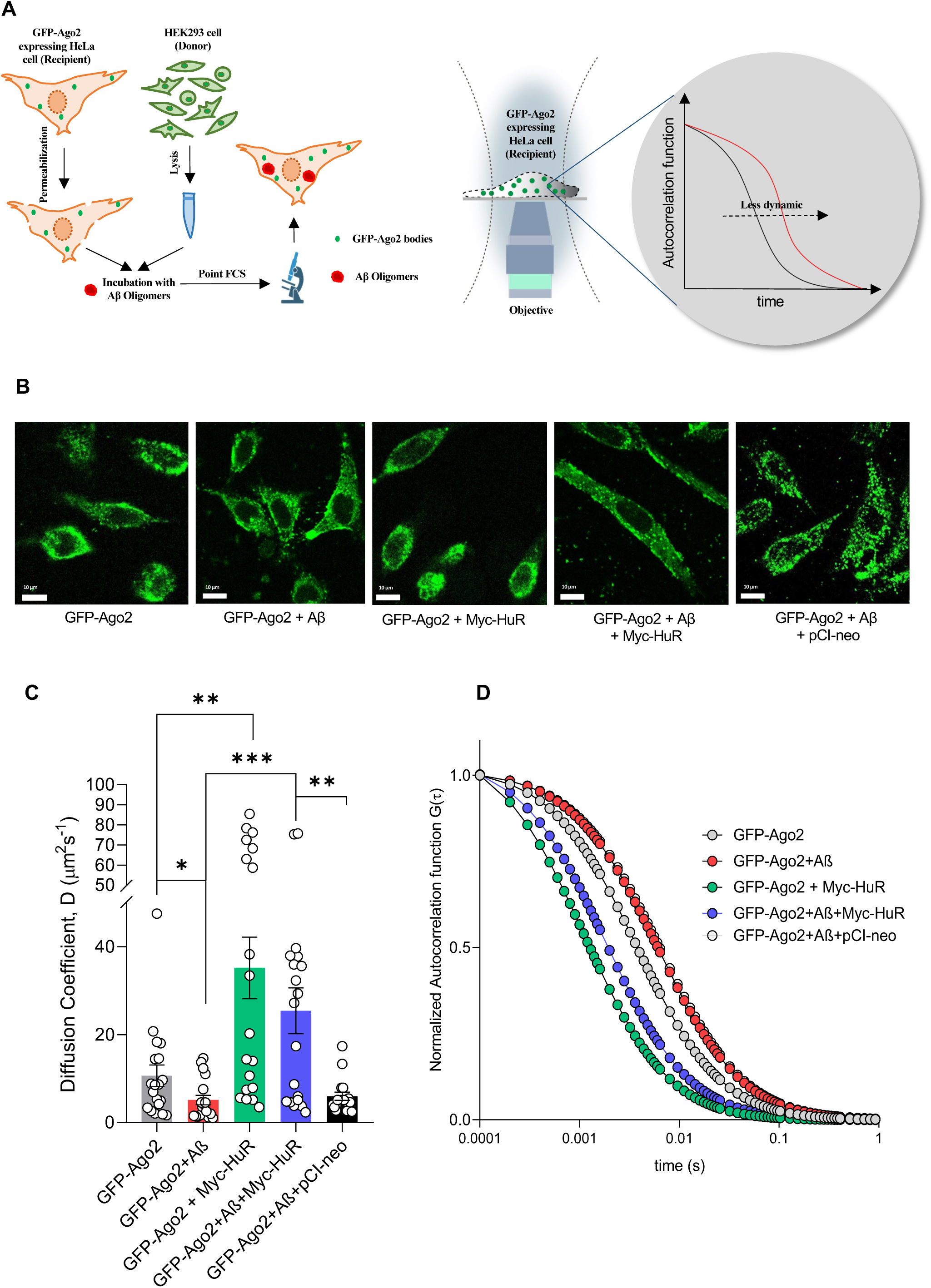
Effect of amyloid beta oligomers on intracellular dynamics and diffusion of phase separated Ago2 bodies. **(A)** A schematic representation of the permeabilization assay using GFP-Ago2 expressing HeLa cells as recipient and HEK293 cells as donor including Aβ treatment for FCS assay (Left panel). A schematic representation of how the point FCS experiment has been done for recipient HeLa cells to obtain the autocorrelation function of GFP-Ago2 protein against time (right panel). **(B)** Live cell images of GFP-Ago2 expressing HeLa cells (recipient) incubated with untransfected HEK293 cells (donor) lysate in absence or presence of 2.5 µM Aβ_(1-42)_ or HEK293 cells (donor) lysate transfected with pCI-neo/ Myc-HuR in absence or presence of 2.5 µM Aβ_(1-42)_. (**C**) Diffusion co-efficient measured by point FCS done on different samples mentioned above (p=0.0417 (between GFP-Ago2 and GFP-Ago2+Aβ groups), p=0.0022 (between GFP-Ago2 and GFP-Ago2+Myc-HuR groups), p=0.0005 (between GFP-Ago2+Aβ and GFP-Ago2+Myc-HuR+Aβ groups), p=0.0071 (between GFP-Ago2+pCI-neo+Aβ and GFP-Ago2+Myc-HuR+Aβ groups), n≥17). **(D)** Normalized autocorrelation function against time was plotted for different samples mentioned above. Fields have been detected in 60X magnification. Scale bar represents 10 µm. Data represents means ± SEMs; ns, non-significant, *p < 0.05, **p < 0.01, ***p < 0.001, ****p < 0.0001. p values were obtained by using two-tailed unpaired Student’s t test.

**Figure 9.**
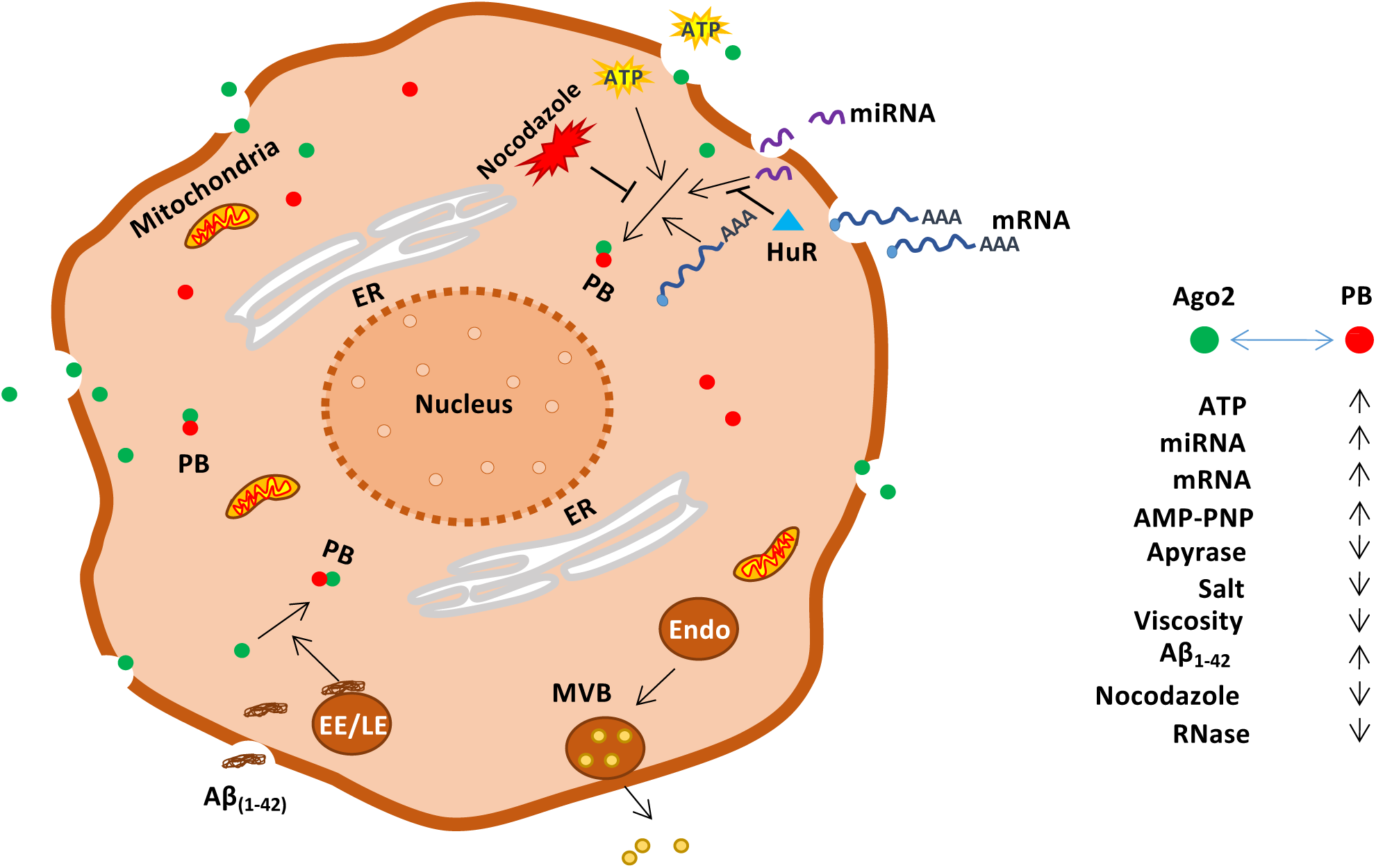
PB phase separation of Ago2 in detergent permeabilized HeLa cells. A schematic representation of the probable factors affecting *in vitro* P-body assembly in recipient HeLa cell. On incubation with FH-Ago2 expressing donor HEK293 cell lysate, permeabilized recipient HeLa cells uptake FH-Ago2 proteins along with other externally supplemented factors present in the lysate through the pores formed by detergent treatment. These internalized FH-Ago2 proteins get targeted to the endogenous P-bodies present in the recipient HeLa cells and this targeting appears to be affected by various factors which are used for manipulating this *in vitro* assay system in various ways for visually studying their effect on the *in vitro* P-body assembly process employing confocal microscopy. Externally added factors like ATP, RNA, miRNA and Aβ_(1-42)_ in the donor lysate showed positive effects on this compartmentalization of internalized FH-Ago2 protein to endogenous P-bodies. On the other hand, negative effects on this phase separation process were observed on Apyrase, RNase and Nocodazole treatments of the lysate which degrade ATP, RNA and microtubule polymerization, respectively. So, these results cumulatively revealed roles of energy, RNA and cytoskeletal structure on in vitro P-body assembly process. The energy dependence of P-body assembly process is further validated by supplementation of AMP-PNP, a non-hydrolysable analogue of ATP, in the lysate which failed to show any impact on the same. Physiochemical properties like salt concentration, viscosity also showed evident effects on this phase separation process causing a gradual decrease in the *in vitro* P-body assembly when their concentrations are gradually increased. ER, Endoplasmic Reticulum;; PB, RNA Processing Bodies; EE, Early Endosome; LE, Late Endosome; Endo, Endosome; MVB, Multi-vesicular Bodies.

Interestingly, we observed a recovery of the arrested dynamics in the presence of Myc-HuR. In this case GFP-Ago2 expressing Hela cells (recipient) were permeabilized and incubated with pCI-neo or Myc-HuR transfected HEK293 (donor) cell lysate in the absence or presence of 2.5 µM Aβ_(1-42)._ Here, pCI-neo was taken as control empty vector plasmid. The diffusion coefficient of GFP-Ago2 was increased significantly (∼ 5 times) in the presence of Myc-HuR (when treated with Aβ_(1-42)_. It is worth mentioning that, even in the absence of Aβ_(1-42)_, Myc-HuR can recover the dynamics of GFP-Ago2. The diffusion coefficient was found to increase ∼ 3.5 times in the absence of Aβ_(1-42)_. A similar shift in auto-correlation function was observed in the case of Myc-HuR also. (Figure 8C and D) In the presence of empty vector pCI-neo, there was no recovery of the reduced dynamics of GFP-Ago2 in the presence and absence of Aβ_(1-42)_.

## Discussion

Liquid-liquid phase separation has emerged in recent years as a vital mechanism explaining formation and function of various membrane less structures present in the cell. Macromolecules undergoing LLPS tend to condense into a dense phase which resembles liquid droplets like structures, co-existing with surrounding dilute phase (13). This phase separation appears to be driven by weak and multivalent interactions between different proteins and nucleic acids, allowing the resulting condensates to exhibit higher protein density but low molecular motion which increases the rate of biochemical reactions in discrete intracellular regions explaining their function in spatio-temporal regulation of many biological activities occurring in the cell (14,26). Two types of macromolecules participate in LLPS-scaffolds and clients. Scaffolds include multivalent proteins and nucleic acid whose multivalency is achieved by the presence of multiple intrinsically disordered regions (IDR) in protein and nucleic acid chains respectively and their higher valency can drive the occurrence of LLPS by enabling the formation of large oligomers and polymers. On the other hand, clients are molecules with lower valency, hence they cannot directly participate in the LLPS process and their recruitment in those condensates is determined by the scaffold molecules (27–29). The multivalent interactions triggering LLPS are again categorized into two varieties-one is the conventional multivalent interaction between protein-protein, protein-RNA, RNA-RNA and the other is weak, multivalent and transient interactions between the IDRs including π-π, dipole-dipole, π-cation, cation-anion interactions (28,30).

In recent studies, LLPS has emerged to be involved in regulating a wide variety of biological activities including adaptive and innate immune signaling, transcription, autophagy, P-body assembly, heterochromatin formation, and miRISC assembly. Any aberration in any of these processes will certainly lead to the development of different disease conditions. In case of neurodegenerative diseases, one of the main hallmarks is protein aggregates. These pathological protein aggregates result from aberrant phase separation process. Disease conditions like Frontotemporal dementia (FD), Alzheimer’s disease (AD), Amyotrophic lateral sclerosis (ALS), Parkinson’s disease (PD) mainly result from an aberrant transition of reversible dynamic LLPS to irreversible aggregation LLPS of fused in sarcoma (FUS), Tau, TAR DNA-binding protein 43 (TDP-43) and α-synuclein proteins respectively. This aberrant phase separation often caused by post-translational modifications (PTM) and disease-associated mutations (31–34). Recently, LLPS also could shed some lights on the relationship between different mutations and the development of cancer, which were unclear for a long period of time. For an example, a mutation in a tumor suppressor protein, Speckle-type POZ protein, can lead to the development of many solid tumors as in prostate, colorectal and gastric cancer. SPOP is a substrate adaptor of a cullin3-RING ubiquitin ligase that mediates the degradation of its substrates including some proto-oncogenic proteins-androgen receptors, death-domain-associated-proteins etc., via the ubiquitin-proteosome system. SPOP consists of a substrate-binding domain MATH and 2 dimerization domains BTB and BACK. A mutation in the MATH domain interferes with the phase separation of SPOP with its substrates into a condensate which happens to be the site of their degradation and this interference ultimately results in the aggregation of these proto-oncogenic proteins leading to the development of cancer condition (35–37). LLPS also plays important role in development by driving the formation and phase separation of germ granules that determine the identity of germ cells during early embryogenesis (38).

However, mechanistic *in vivo* study of phase separation comes with its own challenges. *In vitro* study, on the other side, holds more advantages, as here; phase separation process can be monitored with a definite set of components. LLPS appears to be very sensitive to even slight alterations in the physiochemical conditions like temperature and concentration of protein, RNA as well as salt. Therefore, during *in vitro* study of phase separation, buffer composition, pH and protein concentration should be tightly monitored during assay. Special carefulness is also required during RNA handling to avoid RNase contaminations that can otherwise affect the LLPS outcome. Many biochemical molecules also show remarkable effects on the LLPS process, for an example ATP, which not only acts as a source of energy but also can alter protein solubility. *In vitro* assays can also be easily manipulated to study the effect of different external factors on the phase separation process, which appears to be little difficult while carrying out *in vivo* study (13).

LLPS leads to the formation of a diverse array of membranelles organelles in the cells, one type among them are Cytoplasmic RNA granules which are considered to be the major foci of post-transcriptional gene regulation in eukaryotes(39). Now, among different kind of RNA granules present in the cytoplasm, one important player is ‘Processing bodies’ (P-bodies) which are solely involved in mRNA translation repression or degradation (40). The role of P-bodies or PBs in mRNA turnover becomes further established by the localization of miRNP bound translationally repressed as well as degradation intermediates of target messages in these droplets (41). Therefore, PBs seem to serve as the storage or degradation sites of translationally repressed mRNAs and there occurs a continuous shuttling of messages between polysomes and P-bodies for the temporal and spatial regulation of gene expression in eukaryotic cells (10). The role of PBs in local translation is very prominently observed in neuronal cells as here translationally repressed mRNAs use PBs as a vehicle to reach the distal dendritic region and on reaching the destination they get derepressed, released from the PBs and become actively translated into protein regulating synaptic activity (6). However, there are still many unanswered questions about the mechanistic basis of mRNA compartmentalization and overall assembly of these sub-cellular granules. One among these questions comes from the existence of many probable factors controlling this phase separation process.

A recent study showed a quantitative *in vitro* reconstitution of yeast RNA processing bodies (P-bodies) by combining seven proteins, which are found at high concentrations in PBs, and also RNA at concentrations which are normally seen at the cellular level and it results in the formation of phase separated droplets, quantitatively resembling the protein partition coefficient and dynamics of cellular measurements. Here, only physiological/cellular conditions are used for this *in vitro* reconstitution and the role of RNA in P-body assembly was not completely understood (42). In our study we tried to establish such an *in vitro* system which can be manipulated in many ways mimicking cellular conditions to reveal the role of different internal or externally added factors on the *in vitro* P-body assembly process. We have also explored the role of RNA on P-body formation and dissolution using this assay system.

Cell-resealing assay is considered to be an important tool for localization studies in eukaryotic cells, the basic principle of which is based on the resealing of permeabilized recipient cells with the lysate of donor cells comprising different proteins and RNAs and studying the localization of the internalized factors within the recipient cells (43). We had tried to establish such a kind of a cell-resealing assay here employing HeLa as recipient cell and HEK293 as donor cell, for the localization study of different internalized factors, keeping the morphology and the internal meshwork of the recipient cells closer to the natural condition. By using this assay, we intended to explore the role of different cis and trans acting factors in the process of PB assembly as well as mRNA compartmentalization to these membranelles organelles. Consistent with previously reported data, we have also documented that Ago2 and miRNA binding are sufficient for targeting of the mRNAs to PBs *in vitro*. We have also confirmed the importance of miRNA in PBs compartmentalization of Ago2 which also depends on Ago2 interaction with mRNA. Therefore, this assay may be useful to identify the heterogenicity exits in the PBs that differentially harbor miRNA-Ago2 and other PB specific factors to account for functional diversity of the PBs. With advanced microscopy-based microdissection of this heterogenous PBs from the recipient cells with defined PB structures may be useful in obtaining the biochemical evidence of functional diversity of PBs in terms of mRNA storage or degradation.

The previous *in vivo* data also suggests existence of diversity in PB-targeting of mRNAs as all mRNAs not targeted to PBs. The basis of this section can be studied *in vitro* with this newly developed assay system. Itis interesting to note that amyloid proteins have apositive effect on PB-targeting of Ago2 miRNPs. We suggest that it is a key mechanism that causes the miRNA activity loss in cells expressing amyloid beta. The mechanism of this amyloid beta driven PB compartmentalization of Ago2 is important to negate the effect of amyloid beta on miRNA machinery and on target mRNA expression. Therefore, future use of this assay system may lead us to identification of miRNA regulators to modulate the conditions of diseases like Alzheimer’s Disease caused by amyloidogenic proteins like Ab_(1-42)_ or Tau oligomers.

### Experimental Procedures

#### Cell culture and transfection

HeLa, HEK293 and Huh7 cells were cultured in Dulbecco’s modified Eagle’s medium (DMEM; GIBCO) supplemented with 10% heat-inactivated fetal bovine serum (FBS; GIBCO) and 1% Penicillin-Streptomycin. PC12 cells were cultured in Dulbecco’s modified Eagle’s medium (DMEM; GIBCO) supplemented with 10% heat-inactivated horse serum (HS; GIBCO), 5% heat-inactivated fetal bovine serum (FBS; GIBCO) and 1% Penicillin-Streptomycin.PC12 cells were differentiated with 100 ng/ml NGF (Promega) in differentiation medium containing Dulbecco’s modified Eagle’s medium (DMEM; GIBCO) supplemented with 0.75% heat-inactivated horse serum (HS; GIBCO), 0.25% heat-inactivated fetal bovine serum (FBS; GIBCO). All the transfections were done using Lipofectamine 2000 reagent (Invitrogen) following manufacturers’ protocol. HeLa cells were transfected with 500 ng of plasmids in a 12 well format. HEK293 cells were transfected with 1 µg of plasmids in a 6 well format. Transfections were done for 6 hours for both the cell-lines. Plasmid transfected cells were spitted after 24 hours of the transfection to a larger growing surface.

#### Cell-resealing assay

Recipient HeLa cells (0.3 × 10^6^ cells, on 0.05% gelatin coated 18 mm coverslips) were permeabilized with 80 ng/ml Digitonin (CALBIOCHEM) in 1X PBS (Phosphate Buffered Saline) for 10 minutes at 4 °C and then washed once with 1X PBS to remove excess Digitonin. Donor HEK293 cells (2 × 10^6^ cells) were lysed with lysis solution containing 0.25 M sucrose in DEPC treated water, 10 mM EGTA, 8.4 mM CaCl_2_, 4 mM MgCl_2_, 78 mM KCl, 50 mM HEPES, 1 mM DTT, 10 µg/ml Cycloheximide, 1X EDTA free protease inhibitor cocktail for 30 minutes at 4 °C in mutarotator and sonicated once for 12 seconds. The lysate was collected after centrifugation at 1000 × g for 10 minutes at 4 °C. Permeabilized recipient HeLa cells were treated with donor HEK293 cells lysate for 30 minutes at 37 °C in a humid chamber and then washed once with 1X PBS to remove excess lysate. Lysate treated recipient HeLa cells were then fixed with 4% Paraformaldehyde in 1X PBS. To see the details of reagents used for different treatments and supplementations during the cell-resealing assay, please refer to supplementary table 4.

### Immunofluorescence

Cells on coverslips were fixed using 4% Paraformaldehyde in 1X Phosphate Buffered Saline (PBS) for 30 minutes at room temperature in dark followed by three washes with 1X PBS. Coverslips were then blocked and permeabilized with 1X PBS containing 10% goat serum (Gibco), 20% bovine serum albumin (HiMedia), and 0.1% Triton X-100 (CALBIOCHEM) for 30 minutes at room temperature. Primary antibody, diluted in 1× PBS with 20% BSA, was then added to cells on coverslips and kept overnight in a humid chamber at 4°C. After three washes with 1X PBS for 5 minutes each, Alexa Fluor-labeled secondary antibody diluted in 1X PBS with 20% BSA, was added to cells on coverslips. The coverslips were then placed in a humid chamber and incubated at room temperature for 1 h at dark followed by three washes with 1X PBS for 5 minutes each.The coverslips were then mounted with Vectashield mounting medium with DAPI (4′,6-diamidino-2-phenylindole; Vector).

### Western Blot

The samples were diluted in 5X sample loading buffer (312.5 mM Tris-HCl pH 6.8, 10% SDS, 50% glycerol, 250 mM DTT, 0.5% Bromophenol blue) and heated for 10 minutes at 95°C. Following SDS-polyacrylamide gel electrophoresis, proteins were transferred to PVDF membranes (Milipore). Membranes were blocked in 1X TBS (Tris Buffered Saline) containing 0.1% Tween-20 (TBST) and 3% BSA (HiMedia). Primary antibodies were added in 3% BSA containing TBST for around 16 hours at 4°C. Following overnight incubation with antibody, the membranes were washed at room temperature thrice for 5 minutes each with 1X TBS containing 0.1% Tween-20 (TBST). Washed membranes were incubated at room temperature for 1-1.5 hours with secondary antibodies conjugated with horseradish peroxidase (1:8000 dilution). Excess antibodies were washed three times with TBST at room temperature. Antigen-antibody complexes were detected with West Pico Chemiluminescent, Luminata Forte Western HRP substrate or West Femto Maximum Sensitivity substrates using standard manufacturer’s protocol. Imaging of all western blots was performed using an UVP BioImager 600 system equipped with Vision Works Life Science software (UVP) V6.80.

### In vitro transcription

RL-Con, RL-3XB-let-7a, RL-3XB-miR-122 and RL-5BoxB plasmids were linearized with *HpaI*restriction enzyme (NEB) and *in vitro* transcribed using mMESSAGEmMACHINE Kit (Ambion) and Poly(A) Tailing Kit (Applied Biosystem) following the manufacturers’ protocols to generate m^7^G-capped RNAs. Activity of all *in vitro* transcribed RNAs was validated by Luciferase Assay using Dual Luciferase Assay Kit (Promega).

### Cy3 labelling of oligos

10 µg of oligos against RL sequences were re-suspended in 0.1 M NaHCO3 buffer (pH 8.8), mixed with 30 µl of DMSO containing 1 vial of CY3 monoreactive dye (GE HealthCare) and kept for 24 hours in dark at room temperature (22). Unreacted dye was removed by two rounds of Ethanol precipitations and 3 rounds of purifications using mini Quick Spin Oligo Columns (Roche). The oligos were obtained from Eurogentec having sequences as follows-

RL 1: 5’aT*cacaaagatgatT*ttctttggaaggtT*ca;

RL 2: 5’aT*tagctggaggcagcgT*taccatgcagaaa;

RL 3: 5’aT*agtccagcacgtT*catttgcttgcagT*ga.

### RNA-FISH (Fluorescence in-situ hybridization)

Cells on coverslips were fixed using 4% paraformaldehyde in 1X Phosphate Buffered Saline (PBS) in dark at room temperature for 30 minutes. Coverslips were washed thrice with 1X PBS for 5 minutes each and permeabilized with 70% alcohol overnight at 4 °C. Cells were rehydrated at room temperature for 10 minutes with 2X SSC, 25% Formamide. Cells were hybridized at 37 °C overnight with hybridization solution containing 10% Dextran Sulfate, 2mM Vanadyl Ribonucleoside Complex, 0.02% RNAse-free BSA, 40 µg Salmon Sperm DNA, 2X SSC, 25% Formamide, 30 ng probe in a humid chamber. Cells were washed twice with 2X SSC, 20% Formamide for 15 minutes each at 37 °C and mounted with Vectashield mounting medium with DAPI (4′,6-diamidino-2-phenylindole; Vector) (22).

### Preparation of Amyloid beta

Lyophilized Aβ_(1-42)_ and Aβ_(1-40)_ (American Peptide) was solubilized in 100% 1,1,1,3,3,3, hexafluoro-2-propanol (HFIP) to a final stock concentration of 1 mM. HFIP was removed by evaporation using SpeedVac (Eppendorf). The pellet was solubilized in anhydrous DMSO and sonicated in a bath sonicator for 40 minutes at 37 °C. The final stock solution had a concentration of 5 mM and was stored in −80 °C. Oligomers of both Aβ_(1-42)_ and Aβ_(1-40)_ were prepared by diluting the stock peptide in Phosphate Buffered Saline (PBS) and Sodium Dodecyl Sulphate (SDS) at a concentration of 0.2% to make the concentration of the peptide 400 µM. The solution was incubated at 37 °C for 16 hours. Then, it was further diluted with PBS to a final concentration of 100 µM and incubated at 37 °C for another 16 hours for proper oligomerization before use(17).

### Fluorescence Correlation Spectroscopy

Fluorescence Correlation Spectroscopy (FCS) measures the fluctuations of fluorescence intensity in small (∼femtolitre) confocal volume and provides the diffusion times of the diffusive fluorescent species. Live-cell FCS measurements were carried out using an ISS Alba FFS/FLIM confocal system (Champaign, IL, USA), coupled to a Nikon Ti2U microscope equipped with the Nikon CFI Plan Apo 60X / 1.2NA water immersion objective. The 488-nm picosecond pulsed diode laser was used for the excitation for the FCS measurements. The fluorescence emission was collected using a pair of SPAD (Single Photon Avalanche Detector) detectors with the 50/50 beam splitter and the 530/43-nm band-pass filter. Using two detectors enabled us to determine single colour cross correlation functions to eliminate the artefacts given by the detector after-pulsing. Live cell images of HeLa cells were taken and diffusion was measured in point-FCS mode. The FCS correlation curves were fit to the 3D Gaussian 1-component diffusion model, where the beam waists in the radial and axial dimensions were calibrated using a standard fluorescence dye (Rhodamine 6G, R6G) in water of known diffusion coefficient (2.8 × 10^-6^ cm^2^/sec). The diffusion coefficient of different samples are measured by measuring the diffusion time (τ_D_) and using the relation (44) :

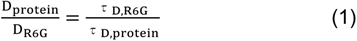

The hydrodynamic radii of each species were computed from their respective diffusion coefficients (D). The FCS data obtained was normalized to free dye following a previously published method (45).

In the 3D Gaussian diffusion model involving a single type of diffusing molecules (the 3D Gaussian 1-component model excluding the contributions of the triplet state), the correlation function G(τ) can be defined by the following equation:

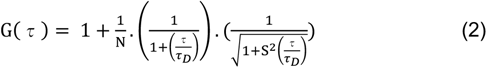

where τ_D_ denotes the diffusion time of the diffusing molecules, N is the average number of molecules within the observation volume, and S is the structural parameter that defines the ratio between the radius and the height. The value of τ_D_ obtained by fitting the correlation function is related to the diffusion coefficient (D) of a molecule by the following equation:

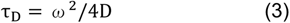

Where ω is the size of the observation volume. From here, the value of the hydrodynamic radius (r_H_) of the protein molecule/ complex/ aggregate can be obtained from D using the Stokes− Einstein formula:

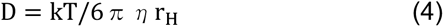

Where η is the viscosity, T is the absolute temperature and k is the Boltzmann constant.

### Image capture and post-capture image processing

FRAP images were captured in Leica confocal microscope SP8 (Leica Microscope Systems; Wetzler, Germany) and all the other imagings were done in Zeiss LSM800 confocal microscope. For FRAP analysis post bleaching recovery was recorded for 1 minute and the images were processed and analysed using LASX Office 1.4.5 27713 software. All other image processing and analysis were done with Imaris7 software developed by BITPLANE AG Scientific software. Pearson’s coefficient of colocalization was calculated using the Coloc plug-in of Imaris7 software. Number and size of individual bodies weremeasured using particle generator of the Surpass plug-in of Imaris7 software.

### Statistical analysis

All graphs and statistical analyses were generated in Prism (v5.00 and v8.00) softwares (GraphPad, San Diego, CA). Nonparametric two-tailed unpaired *t* tests were used for analysis. *P* values of <0.05 were considered statistically significant, and *P* values of >0.05 were not significant. Error bars indicate means ± standard error of the mean (SEMs).

## Supporting Information

Information of plasmids, oligos, antibodies, siRNAs and small RNA are available in Supplementary Table S1 to S4

## Data Availability

All data supporting the findings of this study are available from the corresponding authors upon request.

## Acknowledgements

We thank Witold Filipowicz and Gunter Meister for different constructs used in this study. We thank the Funding body, Dept. of Science and Technology (DST), Govt. of India along with Council for Scientific and Industrial Research (CSIR), for the fellowship to S.R. SNB was supported by The Swarnajayanti Fellowship (DST/SJF/LSA-03/2014-15) from Dept. of Science and Technology, Govt. of India. The work also received support from a High-Risk High Reward Grant (HRR/2016/000093) from Dept. of Science and Technology, Govt. of India and CEFIPRA project grant 6003-J. SNB acknowledge the Start-Up Support Grant of University of Nebraska, USA.

## Author Contributions

S.N.B. and K.M. conceived the idea, designed the experiments, and analyzed the data. S.R. has contributed in design and planning the experiments. S.R. performed the experiments. S.R. and S.R.C performed the FCS experiments. S.R.C. and K.C. analyzed and interpreted the FCS data. S.R., S.B. and Y.C. Standardized the assay system. S.R. and K.M. also wrote the manuscript with S.N.B. and analyzed the data.

